# Cytosolic interaction with RNA-helicase DDX39A titrates viral RNA G-quadruplex mediated α-Synuclein amyloidogenesis

**DOI:** 10.64898/2026.02.24.707865

**Authors:** Aanchal Jain, Shreya Tripathi, Cheshta Agarwal, Anubhab Biswas, Poojitha Sai Potharaju, Aratrika De, Shemin Mansuri, Arindam Mondal, Krishnan H Harshan, Swasti Raychaudhuri

## Abstract

Amyloid aggregates of α-Synuclein are hallmark of Parkinson’s Disease (PD) and related neurodegenerative diseases. α-Synuclein, being an non-canonical RNA-binding protein (RBP), associates with other RBPs within cytosolic RNA-protein granules to modulate mRNA-stability. Conversely, mRNA G-quadruplexes (rG4s) expedite α-Synuclein amyloidogenesis. However, spatiotemporal control on α-Synuclein amyloidogenesis by other RBPs remains unexplored. Here, we report that RNA-dependent cytosolic interaction with DEAD-box RNA-helicase DDX39A decelerates α-Synuclein amyloidogenesis. Viral infections transiently elevate rG4s in cytoplasm. Perturbing interactions between α Synuclein and DDX39A using viral rG4s from H1N1-influenza and SARS-CoV-2 genomes expedites intracellular amyloidogenesis. Conversely, DDX39A overexpression alleviates α-Synuclein amyloidogenesis in mouse primary neurons triggered by SARS-CoV-2 infection. We demonstrate that while DDX39A unwinds viral rG4s to mitigate α-Synuclein sol-gel transition, its reciprocal cooperative phase separation with α-Synuclein enhances the helicase’s rG4-unwinding activity. We propose that accelerated α-Synuclein amyloidogenesis represents a trade-off within this RNA–protein interaction equilibrium, contributing to the viral etiology of PD.

## Introduction

Amyloid aggregates are hallmark of multiple age-related neurodegenerative diseases ^1–4^. Mechanisms including the seeding by pre-formed amyloids, oxidative stress, genetic mutations, deregulation of cellular protein quality control, interactions with gut bacteria and their amyloidogenic proteins; etc. are known to trigger misfolding and aggregation of disease-related amyloidogenic proteins^5–11^. These amyloidogenic proteins localize, form aggregates, and translocate within and between multiple cellular compartments participating in variable interactions and exerting varying degrees of cytotoxicity^12–19^. One common characteristic of multiple amyloidogenic proteins is that these are components of cytosolic or nuclear RNA-protein granules, thus bind to RNA spatiotemporally altering RNA-metabolism^20–24^. On the other hand, intracellular localization modulate their RNA-binding efficiency and liquid-liquid phase separation (LLPS) potential expediting amyloidogenesis ^25–29^. Thus, either way, spatiotemporal interactions between RNA and RNA-binding proteins (RBPs) have functional implications in amyloid-related neurodegeneration.

α-Synuclein is an amyloidogenic protein that shuttles between cell membrane, cytoplasm, nucleus, and other cell organelles^30–35^. Amyloid aggregates of α-Synuclein are associated with many Synucleopathies including Parkinson’s disease (PD)^32,36,37^. Aggregation of α-Synuclein initiates in aging neurons once the endogenous protein undergoes LLPS to nucleate amyloidogenesis^38,39^. α-Synuclein amyloid aggregates are predominantly cytosolic and component of two morphologically and spatiotemporally distinct inclusion bodies (IBs) – perinuclear Lewy bodies (LBs) and axonal Lewy neurites (LNs)^13,40–42^. Spatiotemporal interactions of soluble α-Synuclein, LB-precursors, and mature LBs with RNA, protein, and cell organelles collectively damage cellular homeostasis towards neurodegeneration^13,41–43^. α-Synuclein also associates with multiple RNA-binding proteins (RBPs) in cytosolic processing bodies (P-bodies) to deregulate their function resulting in mRNA-instability^43^. Conversely, cytosolic interactions with mRNA G-quadruplexes accelerates α-Synuclein sol-gel transition and amyloidogenesis^44^. Intriguingly, investigation on α-Synuclein amyloidogenesis via spatiotemporal protein-protein interactions is so far limited to chaperones, trafficking factors etc.^45^. How spatiotemporal RBP-interactions modulate α-Synuclein amyloidogenesis remains unexplored except Tau and TDP-43, which are themselves amyloidogenic^3,46–48^.

Here, we artificially trap α-Synuclein within specific cell compartments in cell culture and mouse primary neuron models to investigate its spatiotemporal interactions and consequent amyloidogenesis. We find that α-Synuclein restricted outside the nucleus interacts with a subset of RBPs that preferentially bind to mRNA-secondary structures. These interactions coincide with the appearance of enlarged processing bodies (P-body) in the cytoplasm. Biogenesis of large perinuclear LB-like inclusions is simultaneously decelerated in these cells. Elevating cellular load of RNA G-quadruplexes (rG4s), including rG4s from the genomes of RNA viruses, perturbs this α-Synuclein-RBP interaction equilibrium and expedites α-Synuclein amyloidogenesis into LB-like inclusions offering a mechanistic explanation of the well-established link between viral infections and an enhanced long-term risk of PD^49–52^. We identify ATP-dependent DEAD-Box Helicase 39A (DDX39A) as an exclusive interactor of α-Synuclein in the cytoplasm. Furthermore, we demonstrate that a tripartite co-condensation equilibrium between α-Synuclein, viral rG4s, and DDX39A calibrates DDX39A rG4-helicase activity as well as the phase-transition landscape of α-Synuclein towards amyloidogenesis.

## Results

### Spatiotemporal confinement modulates α-Synuclein amyloidogenesis

We have recently created a stable and inducible cell culture model in HEK293T cells expressing α-Synuclein variants with EGFP at the C-terminus^13^. Two mutations, A30P and A53T, in α-Synuclein are associated with familial PD^53,54^. Treatment of the cell model expressing an α-Synuclein double mutant containing both A30P and A53T mutations (SNCA(DM)-EGFP) with recombinant α-Synuclein amyloid fibrils (preformed fibrils; PFF) for 6 days resulted in perinuclear LB-like amyloid inclusions (**Fig. 1A, red arrow**). We also created a mouse primary neuron model where endogenous α-Synuclein form perinuclear amyloids upon 10 days PFF-treatment^13^. To confine SNCA(DM)-EGFP in specific cell compartments, we tagged the protein at the C-terminus with nuclear localization signal (NLS) from SV40 large antigen T, nuclear export signal (NES) from MAPK kinase, and the hypervariable domain of K-Ras4B to target the protein to plasma membrane^55–57^. SNCA(DM)-EGFP-NLS was found to be targeted to nucleus preferably **(Fig. 1A)**. SNCA (DM)-EGFP-NES localized predominantly in the cytoplasm while SNCA(DM)-EGFP-KRAS was mostly present at the plasma membrane. Interestingly, western blot of the total cell extracts suggested that membrane-localization prominently reduced the cellular abundance of the protein **(Fig. 1A-B**).

**Fig. 1:**
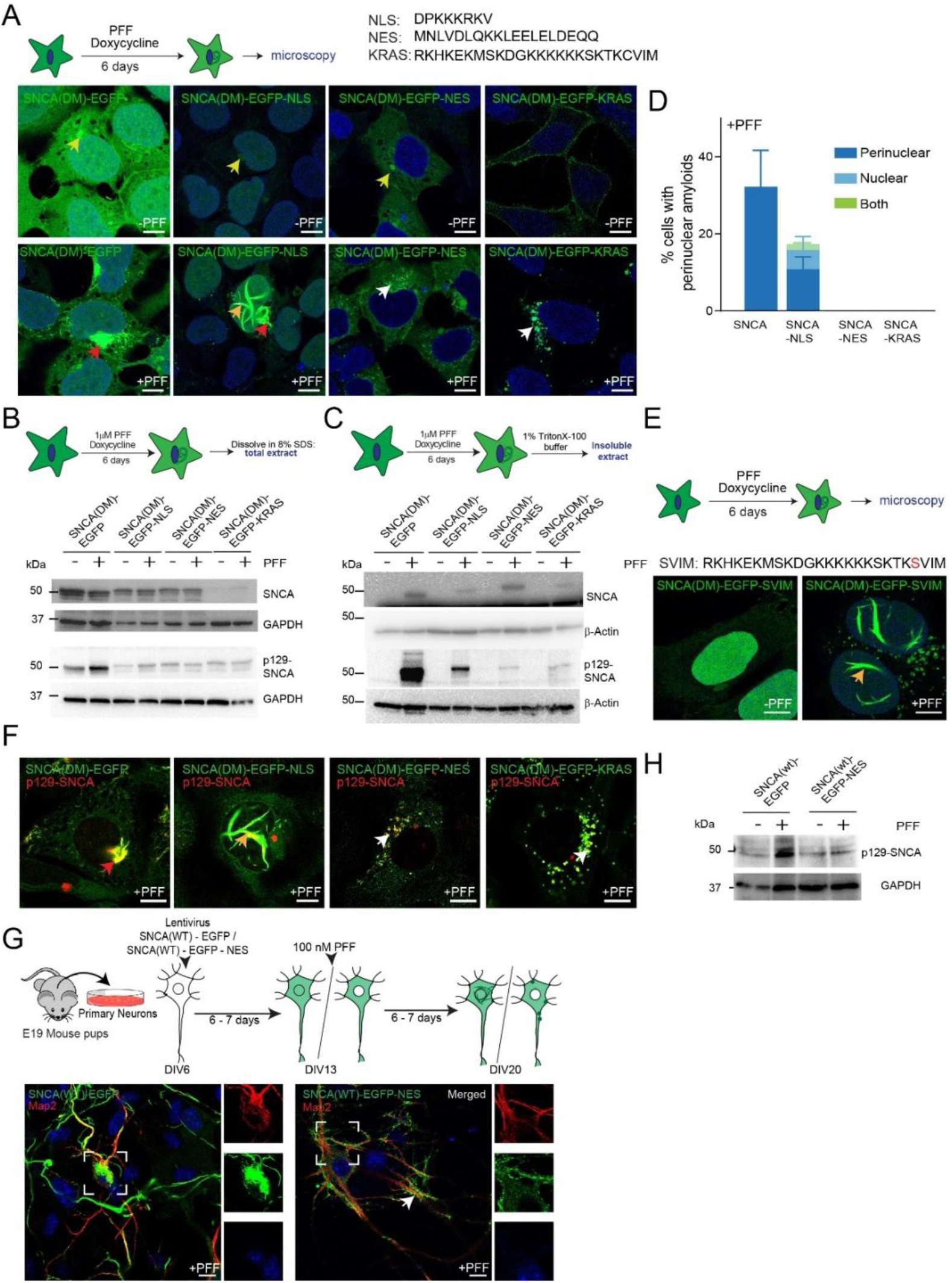
Subcellular location modulates α-Synuclein amyloidogenesis. **A: Top:** Experimental scheme. **Bottom**: HEK293T stable inducible lines expressing EGFP-tagged α-Synuclein (SNCA) and localization variants. Cells were imaged at 6^th^ day in presence or absence of pre formed fibrils (PFF) **B: Top:** Experimental scheme. **Bottom**: Western Blot of total cell lysates probed with SNCA antibodies. SNCA - detecting both phosphorylated and non-phosphorylated α-Synuclein (14H2L1; epitope:117-125) and p129-SNCA - detecting only phosphorylated α-Synuclein (MJF-R13 (8-8)). GAPDH as loading control. **C: Top:** Experimental scheme. **Bottom**: Western Blot of 1% TritonX-100 insoluble extracts probed with SNCA antibodies. SNCA - detecting both phosphorylated and non-phosphorylated α-Synuclein (14H2L1; epitope:117-125) and p129-Syn - detecting only phosphorylated α-Synuclein (MJF-R13 (8-8)). β-actin used as loading control. **D:** HEK293T stable line expressing SNCA(DM)-EGFP and localization variants treated with PFF for 6 days. Number of aggregates containing cells were counted and plotted. Approx. 100 cells were counted in each experiment. N = 3 **E:** Microscopy images of 6^th^ day ± PFF SNCA(DM)-EGFP-SVIM cells. **F:** 6^th^ day PFF-treated SNCA(DM)-EGFP and localization variants were immunostained for p129-SNCA antibody (detecting only phosphorylated α-Synuclein (MJF-R13 (8-8)) and imaged. **G: Top:** Experimental scheme. **Bottom Left**: Mice primary hippocampal neurons at days *in vitro* (DIV) 6 were transduced with lentivirus with SNCA(WT)-EGFP or SNCA(WT)-EGFP-NES, treated with PFF at 13-DIV, fixed and immunostained with anti-MAP2 as neuronal marker at 20-DIV. Insets – single channel zoomed images of boxed regions. **H:** Western Blot of total neuron lysates from **1G** probed with p129-SNCA antibody - detecting only phosphorylated α-Synuclein (MJF-R13 (8-8)). GAPDH as loading control. Error bars indicate standard deviation (SD). Blue staining in microscopy images indicate nucleus (DAPI). Scale bar – 10 μm. Yellow, orange, red and white arrows indicate aggresome, nuclear, perinuclear LB-like amyloid inclusions, and granular IBs, respectively.

We did not observe any aggregates in these cells except aggresomes as revealed by microscopy and western blots of the insoluble fractions^13^ **(Fig. 1A, yellow arrows and C**). PFF-treatment resulted in perinuclear deposition of LB-like filamentous amyloid aggregates in SNCA(DM)-EGFP cells while NLS-tagged version formed both perinuclear and intra-nuclear filamentous aggregates **(Fig. 1A, red/orange arrows and D**). Remarkably, we did not observe filamentous α-Synuclein aggregates in both SNCA(DM)-EGFP-NES and SNCA(DM)-EGFP-KRAS cells rather granular dot-like structures were found surrounding the nucleus upon PFF-treatment **(Fig. 1A, white arrows and D**). More interestingly, mutating the C-terminal cysteine to serine in the KRAS sequence preferentially relocalized the protein (SNCA(DM)-EGFP-SVIM) from plasma membrane to nucleus. In addition to dotted cytoplasmic aggregates, these cells showed formation of intra-nuclear filamentous amyloids upon PFF-treatment (**Fig. 1E**).

Phosphorylation of α-Synuclein at serine 129 (pS129) is a bona fide marker for LB-like amyloid aggregates^41^. The dot-like structures in PFF-treated SNCA(DM)-EGFP-NES and SNCA(DM)-EGFP-KRAS cells were positively immunostained for pS129 **(Fig. 1F and S1A**). Further, the level of pS129-Syn was increased in the insoluble fraction of these cells after PFF-treatment **(Fig. 1C**). The dotted structures in SNCA (DM)-EGFP-NES cells were also positive for Proteastat **(Fig. S1B**), a dye conventionally used to stain amyloid aggregates^38,58^, suggesting these dotted aggregates as precursors of large filamentous LB-like inclusions. We also transduced mouse primary neurons isolated from the hippocampus of embryonic E19 pre-natal C57BL/6 mice with lentivirus expressing either SNCA(wt)-EGFP or SNCA(wt)-EGFP-NES constructs. PFF-treatment resulted in perinuclear LB-like amyloids for SNCA(wt)-EGFP but not for SNCA(wt)-EGFP-NES, which was restricted outside the nucleus (**Fig. 1G-H and S1C**). Instead, only dotted-aggregates were observed at the projections of SNCA(wt)-EGFP-NES neurons (**Fig. 1G, white arrow**). Together, these results suggested that restricting α-Synuclein outside the nucleus decelerated amyloidogenesis towards filamentous LB-like inclusions.

### Spatiotemporal confinement modulates α-Synuclein interactions with RBPs

We hypothesized that difference in α-Synuclein interactions outside or inside the nucleus contributed to its spatiotemporal amyloidogenesis. We immunoprecipitated SNCA(DM)-EGFP variants from our cell lines and performed mass spectrometry to identify the cell compartment specific interactors before PFF-treatment and on 3^rd^ day of PFF incubation when small amyloid filaments begin to appear^13^. Experiments were performed in triplicate with control immunoprecipitations using EGFP with the localization signals. Proteins quantified in at least two replicates were included in the interactome (**Fig. 2A, Table S1**). The unique interactors of SNCA(DM)-EGFP variants were listed first followed by identification of quantitatively enriched interactions by p-value estimation (**Fig. 2A-B, S2A and Table S1**).

**Fig. 2:**
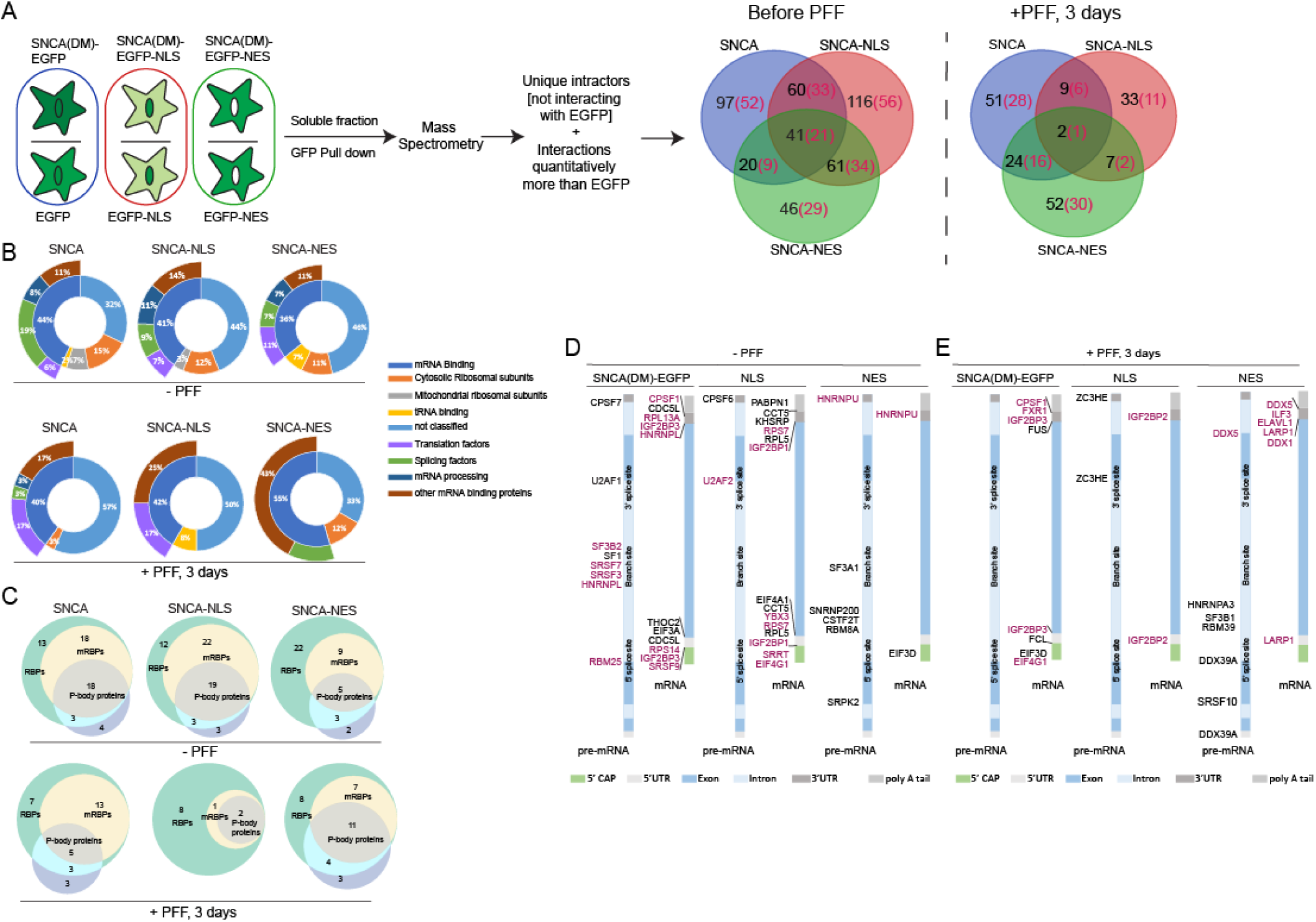
Spatiotemporal confinement modulates α-Synuclein interactions with RBPs. **A:** Experimental scheme and Venn diagram for the immunoprecipitation experiments performed. Venn diagram indicates proteins identified as α-Synuclein interactors in ± PFF treated cells expressing SNCA(DM)-EGFP and localization variants SNCA(DM)-EGFP-NLS and SNCA(DM)-EGFP-NES. Numbers in bracket with red font indicate number of RNA binding proteins (RBPs). **B:** Pie Chart representing distribution of RBPs that are exclusively interacting with α-Synuclein at different cellular localizations categorized into different Gene Ontology (GO) classes. Outer circle represents distribution of mRNA binding proteins. **C:** Distribution of RBPs interacting with α-Synuclein that are exclusively interacting with α-Synuclein at different cellular localizations based on their direct or indirect interactions with P-bodies. Details in **Table S1**. Other mRNA binding proteins are also indicated. **D-E:** Schematic map of pre-mRNA and mRNA structures indicating the possible binding sites of the RBPs within the mRNA schematics. Only subcellular location-specific exclusive interactors of α-Synuclein with literature validated mRNA interactions are mapped. Maroon fonts indicate putative P-body proteins.

Prior to PFF-treatment, the number of interactors for α-Synuclein restricted outside nucleus (SNCA(DM)-EGFP-NES) was less than 50% compared to the interactors observed for α-Synuclein shuttling between cytoplasm and nucleus (SNCA(DM)-EGFP) or predominantly localized to nucleus (SNCA(DM)-EGFP-NLS) (**Fig. 2A**). After 3^rd^ day of PFF-seeding, both SNCA(DM)-EGFP and SNCA(DM)-EGFP-NLS cells showed nearly 50% reduction in number of interactions, suggesting loss of interactions following the onset of filamentous LB-like amyloid formation (**Fig. 2A**). Indeed, amyloid aggregates are known to be less interactive than the soluble amyloidogenic proteins^59^. In contrast, number of interactors in SNCA(DM)-EGFP-NES cells remained unaltered, where only granular aggregates were observed (**Fig. 2A**). This suggested that α-Synuclein persisted in a comparatively amyloidogenic yet soluble state in SNCA(DM)-EGFP-NES cells after 3rd day of PFF-seeding, allowing continued protein interactions.

Gene Ontology (GO) analysis revealed that RNA binding proteins (RBPs) represented more than 50% of the proteins interacting with α-Synuclein variants prior to PFF-treatment irrespective of their subcellular localization (**Fig. 2A and S2B, Table S1**). We further analysed exclusive α-Synuclein interactors at the various subcellular locations. Most of these location specific interactors were found to be mRNA-binding proteins (**Fig. 2B and S2B, Table S1**). Further to that, many of these RBPs were found to be potential components of P-bodies or interactors of the P-body components^60^ (**Fig. 2C**, **Table S1**). This is in accordance to the recent report that α-Synuclein associates with RBPs at the cytosolic P-bodies, modulates the stability of these phase separated RNA-protein granules, and thereby deregulates mRNA metabolism^43^. Interestingly, PFF-treatment reduced interactions with RBPs for SNCA(DM)-EGFP and SNCA(DM)-EGFP-NLS but not for SNCA(DM)-EGFP-NES suggesting a spatiotemporal crosstalk between α-Synuclein, mRNA, and RBPs depending on the intracellular localization of the protein and the stage of amyloidogenesis (**Fig. 2A and S2C**). RBPs predominantly bind to the untranslated regions (UTRs) of the pre-mRNA or mRNA that are known to form secondary structures to regulate splicing, branching, initiation or termination of translation etc.^61,62^. We grouped the α-Synuclein interacting RBPs according to their literature reported binding sites within the pre-mRNA or mRNA sequences (**Fig. 2D-E**). We found that the UTR-binding RBPs prominently reduced as α-Synuclein interactors after 3 days PFF treatment in SNCA(DM)-EGFP and SNCA(DM)-EGFP-NLS cells. Conversely, similar interactions were increased for SNCA(DM)-EGFP-NES (**Fig. 2D-E**). Recently, secondary structures of mRNA have been implicated in expedited amyloidogenesis of α-Synuclein^44^. We hypothesised that spatiotemporal interactions with RBPs that bind to the mRNA secondary structures at the UTRs might not only modulate RNA metabolism as reported earlier^43^ but also amyloidogenesis of α-Synuclein outside the nucleus.

### Cytosolic RNA-protein interactions regulate α-Synuclein amyloidogenesis

P-bodies are phase separated granules that work as the hub of cytosolic mRNA metabolism by facilitating localized RNA-protein interactions^62^. Size and morphology of P-bodies are known to be modulated due to alteration of the stoichiometry of mRNA-metabolizing enzymes within these granules^63–66^. Immunostaining using a canonical P-body component mRNA-decapping enzyme 1A (DCP1A)^67^ revealed 41.66±11.34 tiny puncta (76% ≤ 0.3 and 24% >0.3 µm^2^, median = 0.2 µm^2^) throughout the cytoplasm of SNCA(DM)-EGFP cells (**Fig. 3Ai, white arrows, S3A**). Interestingly, SNCA(DM)-EGFP was found to colocalize with some of these granules (**Fig. 3Ai, insets)**. The number of DCP1A-stained puncta was decreased to 7.119±1.252/ SNCA(DM)-EGFP-NES cells (**Fig. S3A**). The size distribution was also notably different with 42% of puncta ≤ 0.3 and 57% >0.3 µm^2^, median=0.49 µm^2^; suggesting coalescence of multiple P-bodies into enlarged structures when α-Synuclein localization was restricted within cytosol (**Fig. 3Aii, yellow arrows**). Similar observation was reproduced with another P-body component 5’-3’ exoribonuclease XRN1^63^ (**Fig. S3B-C**). Enlarged P-bodies are known to deregulate mRNA-stability by preventing decapping^63^. Thus, emergence of enlarged P-bodies in SNCA(DM)-EGFP-NES cells suggested aggressive modulation of mRNA-metabolism due to cytosolic compartmentalization of α-Synuclein. Interestingly, some of the granular α-Synuclein aggregates observed in 6^th^ day PFF-treated SNCA(DM)-EGFP-NES cells were positive for DCP1A and XRN1 staining suggesting an association between larger P-bodies and these granular amyloids (**Fig. 3Bii-Ci and S3C**). However, the filamentous LB-like amyloids in SNCA(DM)-EGFP cells were not stained by DCP1A antibody (**Fig. 3Bi**). These observations suggested that in addition to altering mRNA-metabolism, RBP-interactions in the proximity of P-bodies in SNCA(DM)-EGFP-NES cells might have arrested α-Synuclein at granular amyloids decelerating its further amyloidogenesis into LB-like aggregates.

**Fig. 3:**
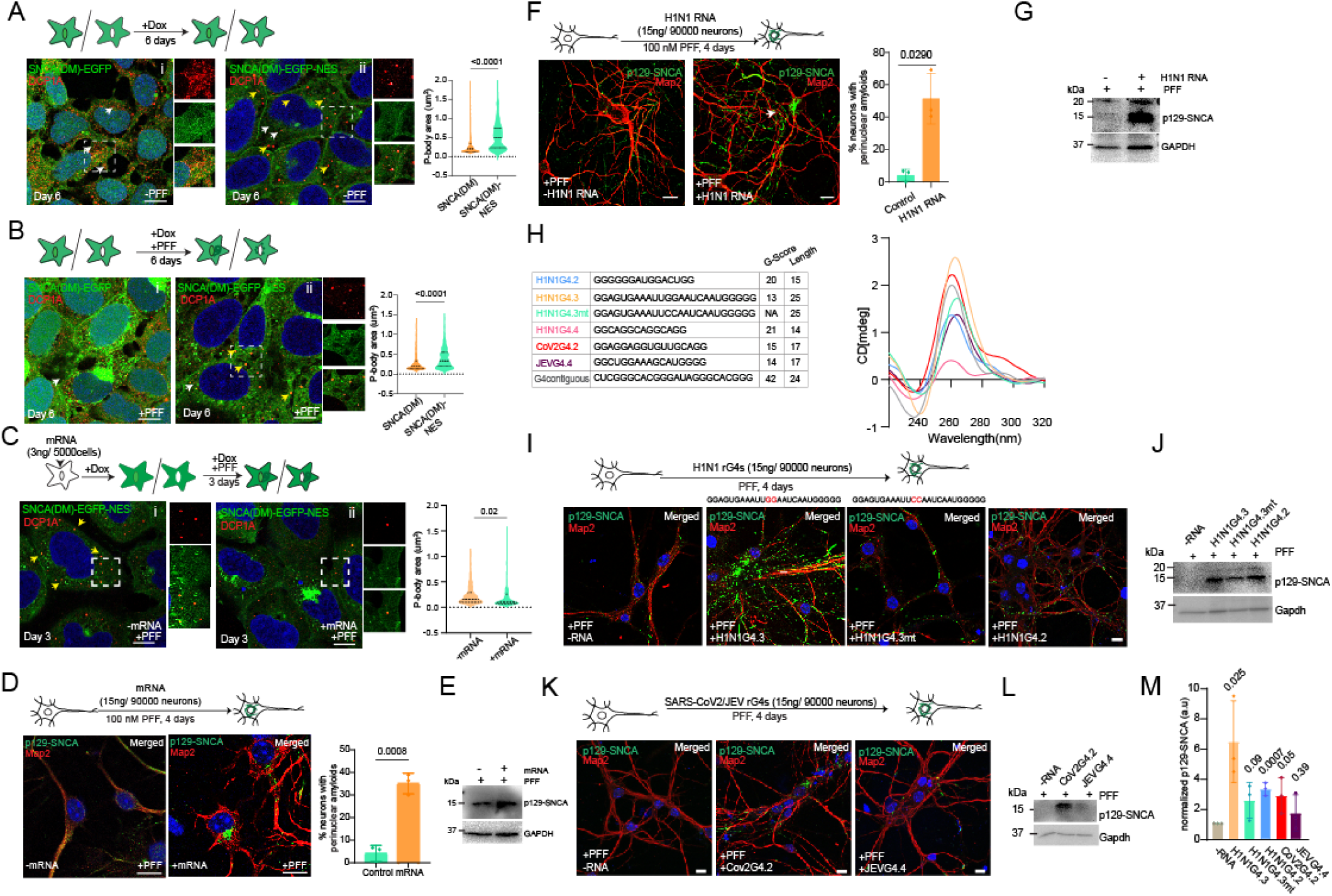
Cytosolic RNA-protein interactions modulate α-Synuclein amyloidogenesis. **A: Top -** Experimental Scheme. **Bottom Left** - Microscopy images of SNCA(DM)-EGFP and SNCA(DM)-EGFP-NES cells in absence of PFF after 6 days immunostained for DCP1A. **Bottom Right** - Violin plot representing size distribution of DCP1A positive P-bodies. Approx. 500 P-bodies were analyzed in each experiment. N = 4. Insets – zoomed single channel images; merged images to show colocalization. White arrow indicates smaller P-bodies and yellow arrows indicate enlarged P-bodies. **B: Top -** Experimental Scheme. **Bottom Left** - Microscopy images showing SNCA(DM)-EGFP and SNCA(DM)-EGFP-NES cells in presence of PFF after 6 days, immunostained for DCP1A. **Bottom Right** - Violin plot representing size distribution of DCP1A positive P-bodies. Approx. 100 P-bodies were analyzed in each experiment. N = 4. Insets – zoomed single channel images; merged image to show colocalization. White arrow indicates smaller P-bodies and yellow arrows indicate enlarged P-bodies. **C: Top -** Experimental Scheme. **Bottom Left** - Microscopy images showing SNCA(DM)-EGFP-NES cells in presence of PFF ±mRNA treatment after 3 days, immunostained for DCP1A. **Bottom Right** - Violin plot representing size distribution of DCP1A positive P-bodies. Approx. 100 P-bodies were analyzed in each experiment. N = 4. Insets – zoomed single channel images; merged images to show colocalization. Yellow arrows indicate enlarged P-bodies. **D: Top -** Experimental Scheme. **Bottom Left** - Mice primary hippocampal neurons at DIV-5 were treated with PFF for 3-DIV ±mRNA, fixed and immunostained with anti-Map2 antibody as neuronal marker and p129-SNCA antibody (detecting only phosphorylated α-Synuclein (MJF-R13 (8-8)) and imaged. **Bottom Right -** Percentage of Map2 positive neurons with perinuclear amyloids. Approx. 20 neurons were counted in each experiment. N = 3. **E:** Western Blot for total cell lysate of mice primary hippocampal neurons in **Fig. 3D** probed with p129-SNCA antibody - detecting only phosphorylated α-Synuclein (MJF-R13 (8-8)). GAPDH as loading control. **F: Top -** Experimental Scheme. **Bottom Left** - Mice primary hippocampal neurons at DIV-5 were treated with PFF for 3-DIV ±H1N1 genomic RNA, fixed and immunostained with anti-Map2 antibody as neuronal marker and p129-SNCA antibody (detecting only phosphorylated α-Synuclein (MJF-R13 (8-8)), and imaged. **Bottom Right** - Percentage of Map2 positive neurons with perinuclear amyloids. Approx. 20 neurons were counted in each experiment. N = 3 **G:** Western Blot of total cell lysate of mice primary hippocampal neurons in **Fig. 3F** probed with p129-SNCA antibody - detecting only phosphorylated α-Synuclein (MJF-R13 (8-8)). GAPDH as loading control. **H: Left -** Table showing RNA G-quadruplex (rG4) sequences from H1N1, SARS-CoV-2 and JEV with G-score as calculated by QRGS mapper and length. G-contiguous from Matsuo et al. used as a positive control sequence^44^. **Right -** Circular dichroism (CD) spectra of rG4s. Color coding in accordance to the table on the left. **I: Top -** Experimental Scheme. **Bottom** - Mice primary hippocampal neurons at DIV-5 were treated with PFF for 3-DIV ±H1N1 rG4s, fixed and immunostained with anti Map2 antibody as neuronal marker and p129-SNCA antibody (detecting only phosphorylated α-Synuclein (MJF-R13 (8-8)), and imaged. **J:** Western Blot of total cell lysate of mice primary hippocampal neurons in **Fig. 3I** probed with p129-SNCA antibody - detecting only phosphorylated α-Synuclein (MJF-R13 (8-8)). GAPDH as loading control. **K: Top -** Experimental Scheme. **Bottom** - Mice primary hippocampal neurons at DIV-5 were treated with PFF for 3-DIV ±SARS-CoV-2 and JEV rG4s, fixed and immunostained with anti Map2 antibody as neuronal marker and p129-SNCA antibody (detecting only phosphorylated α-Synuclein (MJF-R13 (8-8)), and imaged. **L:** Western Blot of total cell lysate of mice primary hippocampal neurons in **Fig. 3K** probed with p129-SNCA antibody - detecting only phosphorylated α-Synuclein (MJF-R13 (8-8)). GAPDH as loading control. **M:** Bar graph representing p129-SNCA levels in mice primary neurons ±viral rG4s as calculated from western blot band intensities normalized by GAPDH. Error bars indicate standard deviation from at least three independent experiments. Blue staining in microscopy images indicate nucleus (DAPI). Scale bar – 10 μm.

To investigate, we doped the cells with excess mRNA extracted from HEK293T cells in order to perturb the α-Synuclein-RBP interaction equilibrium. This resulted in the disappearance of the larger P-body structures in PFF-seeded SNCA(DM)-EGFP-NES cells as revealed by DCP1A staining (**Fig. 3Cii**). Importantly, the disappearance of the enlarged P-bodies was coincidental with the appearance of large LB-like perinuclear aggregates in SNCA(DM)-EGFP-NES cells. Furthermore, addition of mRNA also resulted in expedited amyloidogenesis in SNCA(DM)-EGFP and SNCA(DM)-EGFP-NLS cells (**Fig. S3D**). Appearance of perinuclear LB-like aggregates by 3 days of PFF treatment in the mRNA-doped cells, which otherwise requires PFF-seeding for 6 days^13^, suggested that a disequilibrium of α-Synuclein-RBP interactions by excess mRNA was sufficient to trigger aggressive amyloidogenesis irrespective of intracellular localization of the SNCA(DM)-EGFP variants (**Fig. S3D-E**). Western blots confirmed elevated pS129-Syn in all these mRNA treated cells (**Fig. S3F**). Addition of HEK293T mRNA expedited extensive amyloid formation by endogenous α-Synuclein in mouse primary neurons too (**Fig. 3D**). Large LB-like amyloids surrounded the nucleus already by 4 days of PFF-treatment after mRNA-doping which otherwise requires 10 days to form similar perinuclear inclusions in absence of RNA^13^. Western blots confirmed elevated pS129-Syn in these mRNA treated neurons (**Fig. 3E**). These results are in accordance to the recently published report that RNA G-quadruplexes (rG4s) within cellular mRNA, if present in excess, can accelerate α-Synuclein amyloidogenesis in aging mice brain^44^.

Viral infection can also transiently elevate the cytosolic load of RNA in infected cells^68,69^. Genomes of multiple RNA-viruses contain extensive secondary structures including rG4s^70^. Furthermore, a recent survey using FinnGen project and the UK Biobank (UKB) resources established significant association between negative sense RNA virus H1N1 influenza infection and increased long-term risk of Parkinson’s Disease (PD)^49^. Accordingly, we incubated mouse primary neurons with H1N1 (A/WSN/1933) genomic RNA and observed expedited PFF-mediated amyloidogenesis of endogenous α-Synuclein resulting in perinuclear LB-like pS129-Syn deposits within 4 days (**Fig. 3F**). Western blots confirmed elevated pS129-Syn in the H1N1 genomic RNA treated neurons (**Fig. 3G**).

Next, we predicted potential rG4s within H1N1 (A/WSN/1933) genomic RNA, synthesized the sequences, and confirmed the extent of G-quadruplex formation by circular dichroism (CD) spectroscopy (**Fig. 3H**). The prediction score was highest for H1N1G4.4 sequence followed by the score of H1N1G4.2. These sequence stretches were 15 and 14 nucleotides long, respectively. On the other hand, H1N1G4.3 sequence was 25 nucleotides long with a comparatively low prediction score (**Fig. 3H**). Interestingly, the CD-spectra of H1N1G4.3 showed a characteristic parallel rG4 topology with sharp I_max_ at ∼264 nm and I_min_ at ∼240 nm (**Fig. 3H**). I_max_ at ∼260 nm and I_min_ at ∼245 nm were also observed for H1N1G4.2 but the peak heights were less suggesting a weaker rG4 structure. We observed expedited amyloidogenesis of endogenous α-Synuclein in primary neurons in presence of H1N1G4.3 resulting in perinuclear pS129-Syn+ve amyloid deposits within 4 days (**Fig. 3I**). H1N1G4.2 also triggered α-Synuclein amyloid formation; but to a lesser extent and smaller pS129-Syn+ve amyloid filaments were present at the projections and cell body (**Fig. 3I**). Further, a mutated form of H1N1G4.3 that did not successfully form similarly stable rG4s as the wild type as per CD spectroscopy (**Fig. 3H**), did not effectively trigger aggressive perinuclear amyloidogenesis within the same timeframe (**Fig. 3I**). Like H1N1G4.2, smaller pS129-Syn+ve amyloid filaments were present at the projections of H1N1G4.3^mt^ treated neurons.

Western blots also suggested that pS129-Syn load was comparatively higher in H1N1G4.3 treated neurons than H1N1G4.3^mt^ and H1N1G4.2 treatment (**Fig. 3J**). Further, we tested the possible rG4 sequences from the genomes of positive sense RNA viruses like Japanese Encephalitis Virus (JEV, P20778) and SARS-CoV-2 (Wuhan/WIV04) (**Fig. 3H**). These formed weaker rG4s than H1N1G4.3 as per CD spectroscopy (**Fig. 3H**) and were able to trigger amyloid filaments at the neuronal projections with increased pS129-Syn load than control neurons (**Fig. 3K-L**). Intensity estimation of western blot bands confirmed increased pS129-Syn load in neurons treated with H1N1G4.3 than the other viral rG4s (**Fig. 3M**). Together, these results suggested that like cellular mRNA rG4s, viral rG4s could also expedite intracellular amyloidogenesis of α-Synuclein and thus, provided a robust experimental system to investigate the role of rG4-interacting RBPs in α-Synuclein amyloidogenesis.

### Helicase DDX39A prevents rG4-mediated α-Synuclein amyloidogenesis

Helicases are a set of RBPs that can unwind nucleic acid structures using their ATP-driven molecular motor activity^71,72^. rG4s are inherent in viral genomes and multiple helicases, including the DEAD-box (DDX) helicases, are implicated to unwind viral rG4s to regulate viral replication and propagation^73,74^. We found three DDX helicases DDX1, DDX5 and DDX39A interacting with α-Synuclein (**Fig. 2E**). DDX1 is known to unfold rG4s^75^ and was found as a common interactor of all α-Synuclein variants before the PFF-treatment and of SNCA(DM)-EGFP-NES after PFF-treatment (**Table S1**). DDX5 and DDX39A were exclusive interactors of SNCA(DM)-EGFP-NES after 3 days of PFF treatment (**Fig. 2E, Table S1**). DDX5 is known to unfold DNA G4s at the promoter region of *MYC* gene to activate transcription^76^. Unfolding of G4s by DDX39A is not reported yet. However, this predominantly nuclear helicase translocates to cytoplasm upon positive-sense, single-stranded RNA virus chikungunya (CHIKV) infection, binds to a conserved 5′ sequence element on the viral genome, and prevents viral propagation^74^.

We cloned both DDX5 and DDX39A with mCherry at the C-terminus. DDX5-mCherry was found to be comparatively less abundant than DDX39A-mCherry upon transient transfection in SNCA(DM)-EGFP cells as revealed by western blots (**Fig. 4A, middle panel**). Interaction of both the helicases with α-Synuclein was confirmed by immunoprecipitation by EGFP followed by blotting with mCherry (**Fig. 4B and S4A**). Interestingly, RNase treatment of the cell lysate increased α-Synuclein interaction for DDX39A, but not for DDX5, suggesting an interaction equilibrium between RNA-DDX39A-α-Synuclein (**Fig. 4B and S4A**). Accordingly, DDX39A interaction with SNCA(DM)-EGFP-NES was reduced when we doped the cells with H1N1G4.3 rG4 (**Fig. 4C**). Immunoprecipitation of DDX39A was found to be more in SNCA(DM)-EGFP-NES cells suggesting a cell compartment specificity of the interaction between this helicase and α-Synuclein (**Fig. 4B**). In contrast, DDX5 was found to be similarly interacting with α-Synuclein in both SNCA(DM)-EGFP and SNCA(DM)-EGFP-NES cells (**Fig. S4A**). Importantly, we observed that overexpression of DDX39A, but not DDX5, consistently and significantly reduced the number of perinuclear amyloid containing SNCA(DM)-EGFP cells (**Fig. 4D**) with a notable reduction of pS129-Syn load after PFF treatment (**Fig. 4A, top panel**). DDX39A also colocalized with SNCA(DM)-EGFP perinuclear dots in PFF-treated cells (**Fig. 4D, middle panel**).

**Fig. 4:**
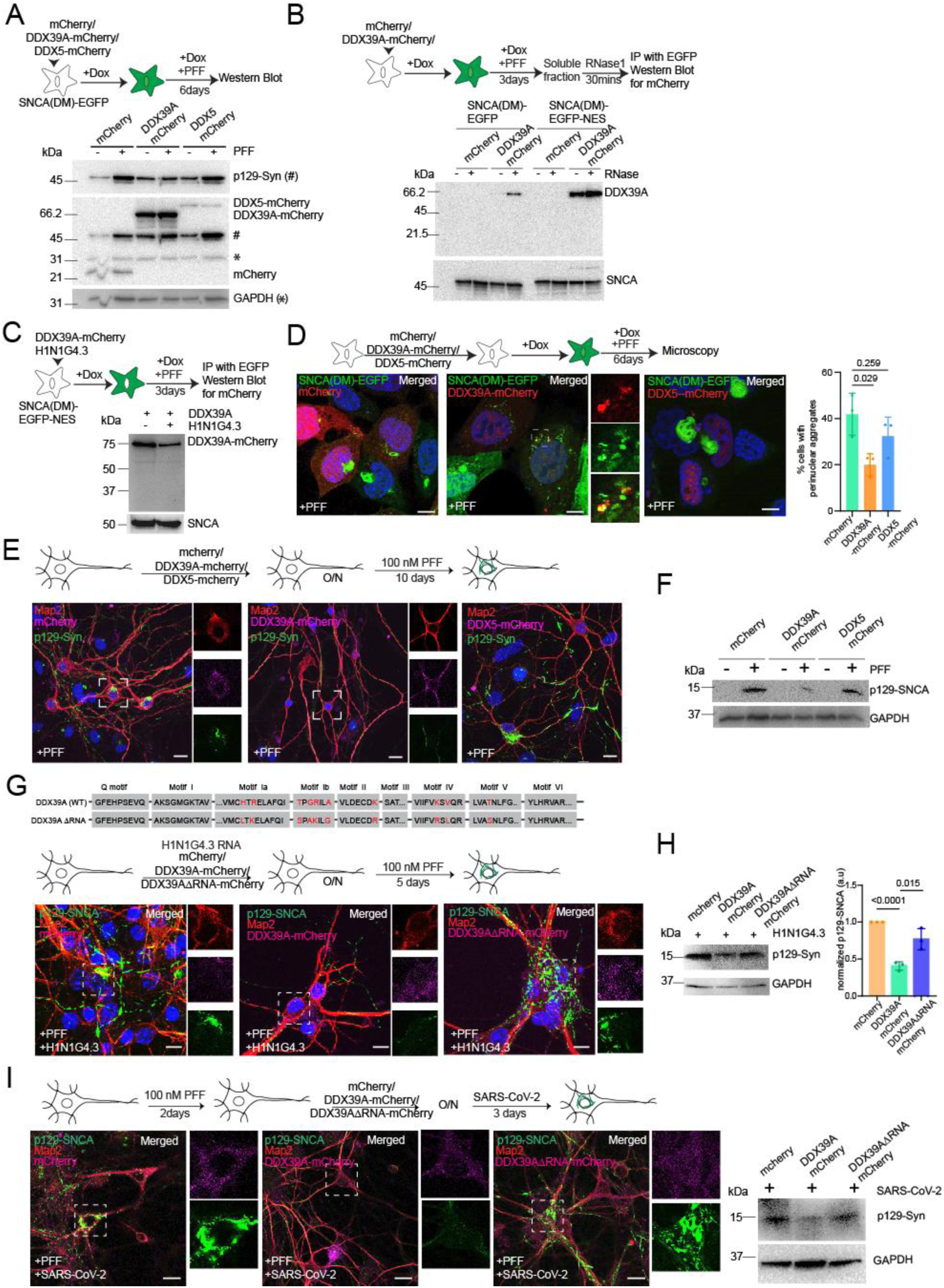
DDX39A alleviates rG4-mediated α-Synuclein amyloidogenesis. **A: Top -** Experimental scheme. **Bottom** - Western blot of total fractions of SNCA(DM)-EGFP cells transfected with mCherry, DDX39A-mCherry and DDX5-mCherry ±PFF for 6 days. Blots were probed with p129-SNCA antibody - detecting only phosphorylated α-Synuclein (#, MJF-R13 (8-8)) and mCherry antibody. GAPDH as loading control (*). Bands for full length DDX5-mCherry and DDX39A-mCherry indicated. **B: Top -** Experimental scheme. **Bottom** – SNCA(DM)-EGFP and SNCA(DM)-EGFP-NES cells were transfected with mCherry and DDX39A-mCherry and treated with PFF for 3 days. Immunoprecipitation (IP) for GFP was performed with soluble fractions ±RNase1 (30 min) and probed with SNCA antibody - detecting both phosphorylated and non-phosphorylated α-Synuclein (14H2L1; epitope:117-125) and mCherry antibody. **C: Top -** Experimental scheme. **Bottom** - SNCA(DM)-EGFP-NES cells ± H1N1G4.3 rG4 was transfected with mCherry and DDX39A-mCherry and treated with PFF for 3 days. Immunoprecipitation (IP) for EGFP was performed with 1% TritonX-100 soluble fractions and probed with SNCA antibody - detecting both phosphorylated and non-phosphorylated α-Synuclein (14H2L1; epitope:117-125) and mCherry antibody. **D: Top -** Experimental scheme. **Bottom left** – Microscopy images of SNCA(DM)-EGFP cells transfected with mCherry, DDX39A-mCherry and DDX5-mCherry ±PFF for 6 days. Insets – zoomed single channel images, merged image indicates colocalization. **Bottom right** – percentage of cells with perinuclear aggregates. Approx. 100 cells were counted in each experiment. N = 3 **E: Top -** Experimental scheme. **Bottom** - Microscopy images of mice primary hippocampal neurons at DIV-5 transfected with mCherry, DDX39A-mCherry and DDX5-mCherry +PFF for 10 days, immunostained with anti-Map2 antibody as neuronal marker and p129-SNCA antibody (detecting only phosphorylated α-Synuclein (MJF-R13 (8-8)). Insets – zoomed single channel images. **F:** Western Blot for total lysate of mice primary hippocampal neurons in **Fig. 4E** probed with p129-SNCA antibody - detecting only phosphorylated α-Synuclein (MJF-R13 (8-8)). GAPDH as loading control. **G: Top -** Sequence map showing putative RNA binding motifs of DDX39A(WT) protein and RNA binding mutations in DDX39AΔRNA protein^77^ and Experimental scheme. **Bottom** - Microscopy images of mice primary hippocampal neurons at DIV-5 transfected with mCherry, DDX39A-mCherry and DDX39ΔRNA-mCherry +PFF/±H1N1 rG4s, immunostained with anti-Map2 antibody as neuronal marker and p129-SNCA antibody (detecting only phosphorylated α-Synuclein (MJF-R13 (8-8)). Insets – zoomed single channel images. **H: Left** - Western Blot for total lysate of mice primary hippocampal neurons in **Fig. 4G** probed with p129-SNCA antibody - detecting only phosphorylated α-Synuclein (MJF-R13 (8-8)). GAPDH as loading control. **Right** - Bar graph showing p129-SNCA levels as calculated from band intensities from western blots normalized by GAPDH. N=3, Student t-test, Error bars represent SD. **I: Top -** Experimental scheme. **Bottom** - Mice primary hippocampal neurons at DIV-5 were treated with PFF. After 2 DIV neurons were transfected with mCherry, DDX39A-mCherry and DDX39ΔRNA-mCherry along with SARS-CoV-2 infection. Neurons were fixed after 3 days of infection and immunostained with anti Map2 antibody as neuronal marker and p129-SNCA antibody (detecting only phosphorylated α-Synuclein (MJF-R13 (8-8)) and imaged. Insets – zoomed single channel images. **J:** Western Blot for total lysate of mice primary hippocampal neurons in **Fig. 4I** probed with p129-SNCA antibody - detecting only phosphorylated α-Synuclein (MJF-R13 (8-8)). GAPDH as loading control. Error bars indicate SD from at least three independent experiments. Blue staining in microscopy images indicate nucleus (DAPI). Scale bar – 10 μm. O/N - overnight

Immunostaining with pS129-Syn antibody suggested that overexpression of DDX39A reduced the PFF-mediated amyloidosis of endogenous α-Synuclein in mouse primary neurons as well (**Fig. 4E**). Microscopy and western blots further indicated that DDX5 was not as efficient as DDX39A in decelerating PFF-mediated α-Synuclein amyloidogenesis in neurons (**Fig. 4E-F**). We further found that DDX39A was capable to prevent H1N1 genomic RNA mediated (**Fig. S4B**) and H1N1G4.3 mediated expedited amyloidogenesis of endogenous α-Synuclein in mouse primary neurons (**Fig. 4G-H**). To validate further, we created a RNA-binding mutant of DDX39A as described by Shi et al.^77^ H1N1G4.3 mediated α-Synuclein amyloidogenesis remained unperturbed despite overexpression of the mutant in primary neurons suggesting RNA-binding activity as a requirement for the anti-amyloid activity of DDX39A (**Fig. 4G-H**). Since SARS-CoV-2 genomic rG4s were capable to trigger α-Synuclein amyloidogenesis (**Fig. 3I-K**), we next infected primary neurons obtained from K18-hACE2 mice pups with B1.1.8 variant of SARS-CoV-2^78,79^. Infection was confirmed by viral titre estimation and immunostaining of the neurons for viral dsRNA (**Fig. S4C**). PFF-mediated perinuclear α-Synuclein amyloids were observed in the infected neurons after 3 days of infection with an increase of pS129-Syn load in western blot (**Fig. S4C-D**). Intriguingly, DDX39A reduced α-Synuclein amyloidogenesis in these virus-infected neurons while its RNA-binding mutant was not effective (**Fig. 4I**). Taken together, these results suggested that a tripartite cytosolic interaction equilibrium between α-Synuclein, viral rG4s, and RNA helicase DDX39A calibrated the kinetics of intracellular amyloidogenesis of α-Synuclein.

### DDX39A helicase activity regulates rG4-mediated phase transition and amyloidogenesis of α-Synuclein

DDX39A, also known as UAP56-related helicase, 49 kDa (URH49), has been reported to be implicated in the maintenance of genome stability, mRNA export, innate immunity, cancer metabolism, and antiviral activities^74,77,80–82^. This ATP dependent RNA helicase has been shown to unwind RNA-DNA hybrids^81,83^, however, its rG4-unwinding function was never tested. Further to that, DDX39A contains an N-terminal 34 residue unstructured stretch^84^. Electrostatic coacervation with unstructured regions of other proteins has been shown to modulate liquid-liquid phase separation (LLPS) of α-Synuclein^48^. We observed that over-expression of DDX39A could alleviate PFF-mediated amyloid formation by α-Synuclein both in presence or absence of RNA-doping (**Fig. 4E-H**). This suggested that in absence of RNA-doping, DDX39A could either unwind endogenous rG4s to mitigate cellular RNA-driven LLPS of α-Synuclein or modulate α-Synuclein LLPS through its unstructured N-terminus.

To confirm, we purified recombinant DDX39A using His-tag, which was found to bind to H1N1G4.3 rG4 in concentration dependent manner as revealed by electrophoretic mobility shift assay (EMSA) (**Fig. S5A-B**). Importantly, the RNA-binding capacity of DDX39A was ATP-independent (**Fig. S5B**). DDX39A also exhibited an ATP-dependent RNA-helicase activity in rG4-unwinding assays. H1N1G4.3 and CoV2-G4.2 rG4 sequences were incubated with purified DDX39A (0.5 µM) in presence and absence of ATP. Bands indicative of RNA secondary structures disappeared on the native PAGE when the rG4s were incubated with DDX39A and ATP suggesting loss of intermolecular RNA-RNA interactions (**Fig. 5A**). We also performed fluorescence resonance energy transfer (FRET) assay^85^ in which H1N1G4.3 rG4 (40 nM) was flanked by 6-fluorescein (6-FAM) at the 3’-end and black hole-1 quencher on the 5’-end. ATP-driven increase of 6-FAM fluorescence in presence of DDX39A (20 nM) confirmed rG4-unwinding activity of the helicase (**Fig. 5B**).

**Fig. 5:**
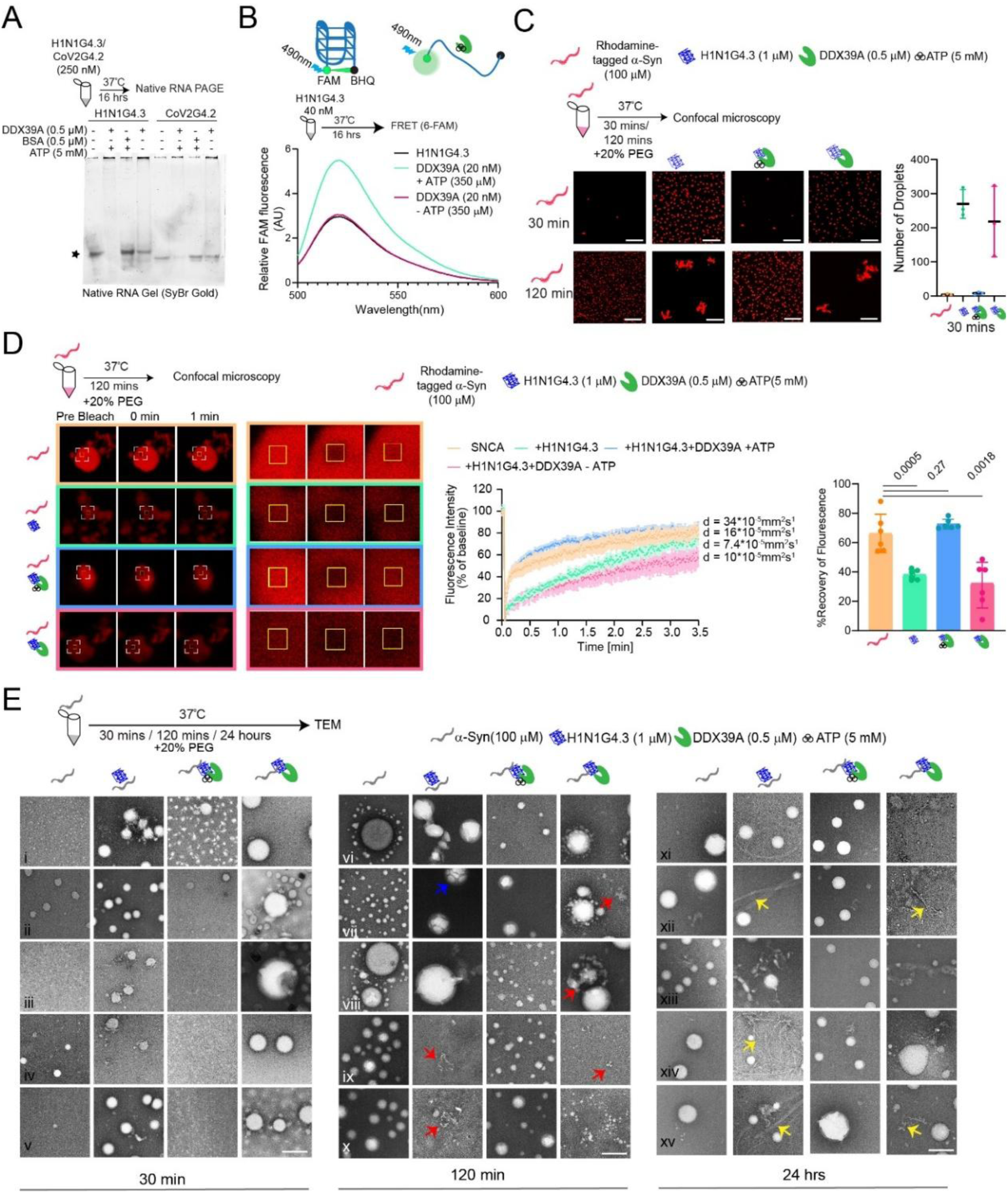
ATP dependent helicase activity of DDX39A prevents α-Synuclein sol-gel transition. **A: Top - Experimental scheme. Bottom** – secondary structures of H1N1G4.3 and CoV2G4.2 rG4s (250 nm each) in presence of 0.5 µM DDX39A ±5 mM ATP as visualized on native RNA gel. 0.5 µM of Bovine serum albumin was used as control. Star indicates H1N1G4.3 G4-quadruplex band. **B: Top -** Experimental scheme. **Bottom –** 6-FAM signal of 40 nM of H1N1G4.3 rG4 in presence of 20 nM DDX39A ±350 μM ATP as observed in FRET assay. **C-E: Top -** Experimental scheme. **Bottom**- *in vitro* LLPS of purified recombinant rhodamine-tagged human α-Synuclein (100 µM) in the presence of 20% PEG. Reactions were set as conditions indicated in figures. **C. left**: Fluorescence microscopy. **Right**: number of droplets per field. 2 fields per repeat were counted, N=3. **D**. FRAP assay. **Left:** fields of microscopy, beaching zones are boxed. **Middle**: FRAP measurements of α-Synuclein droplets. d = diffusion coefficient. The data represent mean ± s.e.m. **Right**: Percent recovery of fluorescence. N = 6 droplets from 3 independent experiments. Error bars indicate SD. **E.** Transmission electron microscopy (TEM) images. 5 fields per experimental condition. Color-codes of arrows indicate different phase separated states as described in text. Scale bar – 10 μm for **5C**, Scale bar – 1 μm for **5D**, Scale bar – 200 nm for **5E**

Next, we setup an *in vitro* experiment to investigate α-Synuclein LLPS in presence of molecular crowding agent polyethylene glycol (PEG)^38^. When α-Synuclein (100 µM) was incubated with 20% PEG at 37°C for 30 minutes, presence of H1N1G4.3 rG4 (1 µM) resulted in approximately ∼250 condensates per field, compared to just 4-5 droplets in the control, implying that rG4 acted as a scaffold for α-Synuclein amyloid formation (**Fig. 5C**). In presence of DDX39A (0.5 µM) and ATP, droplet numbers remained low (around 6-7), while removing ATP negated the inhibitory effect DDX39A on rG4-mediated condensate formation (**Fig. 5C**). When the reactions were prolonged for 120 min, α-Synuclein alone produced ∼100 spherical condensates per field. With H1N1G4.3, few droplets further transitioned into irregularly shaped aggregates due to clustering of the smaller condensates (**Fig. 5C and S5C**). Importantly, fusion of the smaller condensates into the irregular aggregates was restricted in presence of DDX39A and ATP (**Fig. 5C and S5C**). Both the individual droplets and irregular aggregates were positive for proteostat staining indicating amyloid-like properties (**Fig. S5C**).

In contrast, when α-Synuclein was incubated in absence of rG4s, we observed 4-5 liquid droplets of rhodamine-tagged α-Synuclein (100 µM) per field within 30 min of incubation at 37°C (**Fig. S5D**). The number of α-Synuclein droplets was increased to ∼10-12 droplets per field within 30 min from 4-5 in control when we added 0.5 µM DDX39A, with or without ATP, in the reaction mix. The number of droplets was expectedly much more after 2 hr incubation with emergence of few coalesced droplets in samples with DDX39A (**Fig. S5D**). This suggested that the unstructured region of DDX39A, or the full-length protein, did not reduce rather facilitated α-Synuclein LLPS, however, not to the extent of H1N1G4.3 rG4. Together, these observations suggested that H1N1G4.3 first promoted the initial scaffolding of α-Synuclein into phase-separated droplets and further advanced their subsequent fusion into aggregates. Whereas DDX39A, at 1:200 molar ratio with α-Synuclein, remodelled and mitigated the rG4-driven phase transition of α-Synuclein through its ATP-dependent rG4-helicase activity,

To characterize the dynamics of α-Synuclein inside the droplets, we conducted fluorescence recovery after photobleaching (FRAP) experiments at 120 min of the reaction. Under control conditions, α-Synuclein droplets exhibited a rapid fluorescence recovery with a half-time (t₁/₂) of 53.38 seconds and ∼60% recovery, indicating a dynamic, liquid-like state (**Fig. 5D**). Conversely, upon addition of H1N1G4.3, both the recovery rate and extent dropped significantly (t₁/₂ = 115.2 seconds; ∼30% recovery), suggesting a transition toward a comparatively gel-like or solid-like state within the fused droplets (**Fig. 5D**). Addition of DDX39A along with ATP restored fluorescence recovery (t₁/₂ = 25.04 seconds; ∼70% recovery), implying enhanced protein mobility and reversal of the sol-gel transition (**Fig. 5D**). In contrast, this recovery was absent without ATP, highlighting that ATP hydrolysis was essential for the rG4-unwinding activity of DDX39A, which prevented the maturation of α-Synuclein condensates into less dynamic, gel-like states.

Next, we performed transmission electron microscopy (TEM) to obtain more information on the phase transition states of α-Synuclein. After 30 minutes of incubation with PEG at 37°C, α-Synuclein alone exhibited minimal condensate formation. In contrast, the addition of H1N1G4.3 led to the emergence of droplets with heterogeneous sizes (**Fig. 5Ei-v**). Such droplets were absent when DDX39A and ATP were included into the reaction, however, withdrawal of ATP restored droplet formation (**Fig. 5Ei-v**). Notably, in the absence of ATP, reactions containing DDX39A generated larger droplets than those observed with H1N1G4.3 alone (**Fig. 5Ei-v**).

Phase separation was observed for α-Synuclein alone when the reactions were extended for120 min. We observed droplets of heterogeneous sizes (**Fig. 5Evi-x**). At this point, H1N1G4.3-induced condensates appeared even larger than those formed with the rG4 at 30 min (**Fig. 5Evi-viii**). These larger condensates frequently exhibited internal partitioning into multiple irregular sub-droplets, consistent with a sol–gel phase transition (**Fig. 5Evi-vii, blue arrows**). In some fields, droplets were observed depositing onto irregular structures, suggesting ongoing reorganization of phase-separated α-Synuclein toward aggregation (**Fig. 5Eix-x, red arrows**). Similar irregular structures and enlarged droplets were also observed with DDX39A in the absence of ATP. However, in the presence of ATP, DDX39A decelerated H1N1G4.3-induced phase transition, as indicated by the appearance of only smaller, more uniform droplets (**Fig. 5Eix-x**). After 24 hours, reactions containing DDX39A and ATP still displayed only droplets, whereas filamentous amyloids were detected in many fields with H1N1G4.3 rG4 (**Fig. 5Exi-xv, yellow arrows**).

Collectively, these findings suggested that although DDX39A could modestly promote α-Synuclein LLPS in the absence of rG4s, it counteracted rG4-driven acceleration of α-Synuclein phase transition in an ATP-dependent manner by unwinding the RNA secondary structures.

### α-Synuclein facilitates DDX39A phase transition to enhance its helicase activity

Interestingly, DDX39A overexpression did not only reduce α-Synuclein amyloidogenesis in SARS-CoV-2 (B1.1.8)^86^ infected neurons (**Fig. 4I**) but also significantly reduced the viral titre (**Fig. 6A**). Analysis of genome sequence of this SARS-CoV-2 variant revealed multiple potent rG4s within and beyond the coding regions (**Fig. S6A**). Although SARS-CoV-2 variants are continuously evolving, rG4s are inherent to the viral genome and implicated in viral replication and translation processes^87,88^. We posited that overexpressed DDX39A uncoiled these rG4s to inhibit viral replication in our experiments. Interestingly, over-expressed DDX39A was found to colocalize with α-Synuclein in phase separated cytosolic granules (**Fig. 4D**). Multiple DDX helicases have been reported to phase separate and DDX39A itself was found in amyloid bodies in stressed cells^89,90^. Further to that, phase separation of multiple proteins has also been correlated with their enhanced enzymatic activity^91,92^.

**Fig. 6:**
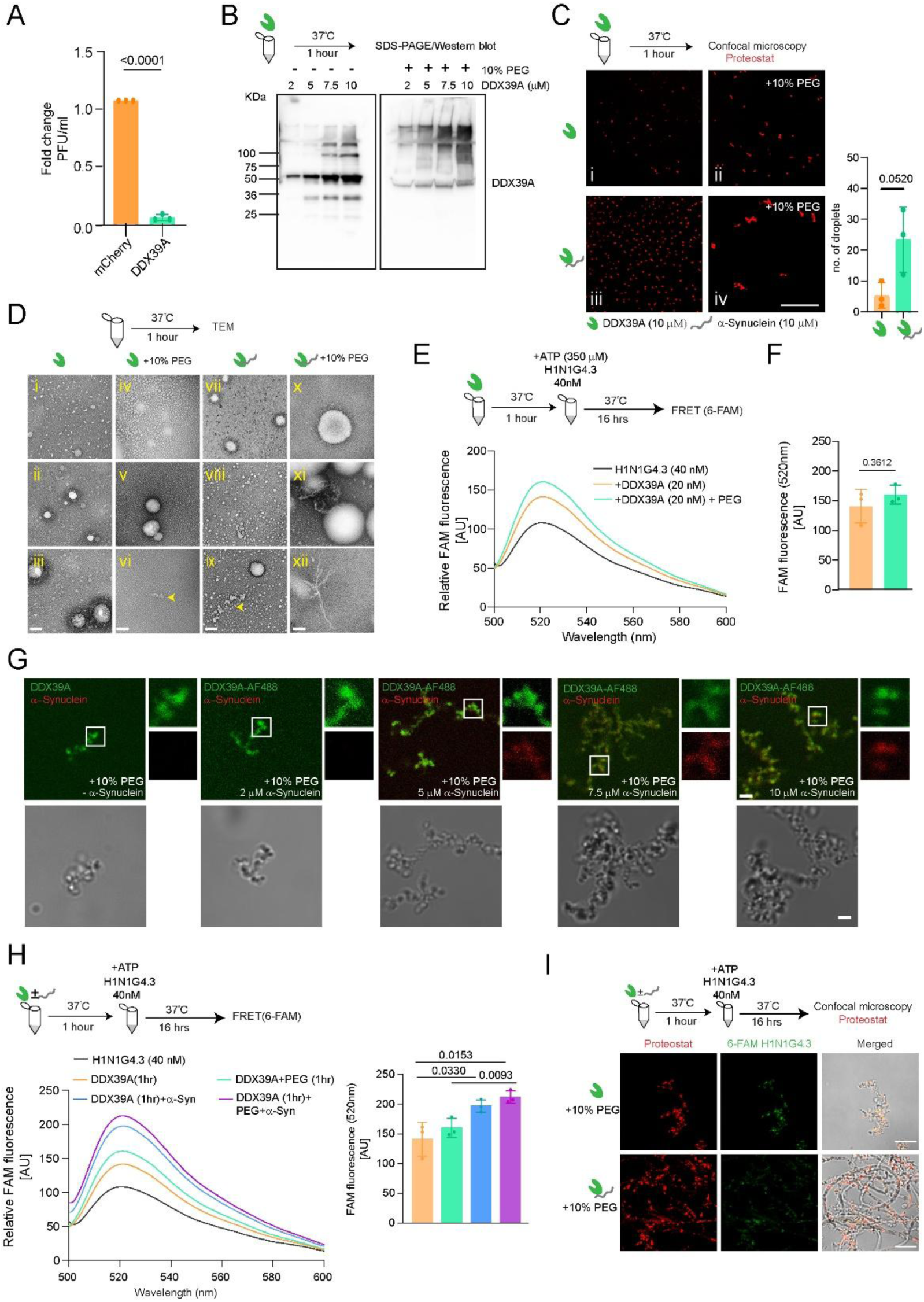
α-Synuclein facilitates DDX39A phase transition to enhance its helicase activity. **A:** Virus titration of supernatant collected after 72 hrs of SARS-CoV-2 infected mice primary hippocampal neurons. N=3, Error bars represent standard deviation, Student t-test. **B: Top -** Experimental scheme. **Bottom –** Purified DDX39A was kept in increasing concentration for 1 hour in absence and presence of 10% PEG. Western blot for the samples was probed with anti-DDX39A antibody. **C: Top -** Experimental scheme. **Bottom** - Microscopy images showing *in vitro* LLPS of 10 µM purified DDX39A in the absence and presence of 10% PEG with and without α-Synuclein after proteostat staining. Scale bar – 10 μm. **Bottom right**: number of droplets in Ci and iii. 2 fields per repeat were counted, N=3. Scale bar – 10 μm. **D: Top -** Experimental scheme. **Bottom** – TEM images showing *in vitro* LLPS of 10 µM purified DDX39A in the absence and presence of 10% PEG with and without α-Synuclein. Scale bar – 100 nm. **E: Top -** Experimental scheme. **Bottom** – *in vitro* LLPS of 10 µM purified DDX39A in the absence and presence of 10% PEG was set up. For FRET, 6-FAM signal of 40 nM of H1N1G4.3 rG4 in presence of LLPS reaction mixture equivalent to 20 nM DDX39A after phase separation ±350 μM ATP was recorded. **F:** 6-FAM signal at 520nm wavelength as observed in FRET assay in **6E** is plotted. N=3, Error bars represent standard deviation, Student t-test. **G:** Microscopy images showing *in vitro* LLPS and co-condesation of purified DDX39A (10µM) labelled with Alexafluor-488 in the presence of 10% PEG with increasing concentration of rhodamine labelled α-Synuclein. Scale bar – 1 μm. **H: Top -** Experimental scheme. **Bottom Left** – *in vitro* LLPS of 10 µM purified DDX39A in the absence and presence of 10% PEG ± α-Synuclein. For FRET, 6-FAM signal of 40 nM of H1N1G4.3 rG4 in presence of LLPS reaction mixture equivalent to 20 nM DDX39A after phase separation ±350 mM ATP was recorded. **Bottom Right** - 6-FAM signal at 520nm as observed in FRET assay plotted. N=3, Error bars represent standard deviation, Student t-test. **I:** Microscopy images showing *in vitro* LLPS of purified DDX39A (10µM) in the presence of 10% PEG without and with α-Synuclein 16 hrs after the addition of 6-FAM labelled H1N1G4.3. Scale bar – 1 μm.

Accordingly, we setup an *in vitro* LLPS reaction for DDX39A. When the reaction-mixes were run on SDS-PAGE followed by western blot, increasing abundance of high molecular weight bands indicated the concentration dependence of DDX39A phase separation (**Fig. 6B**). We observed phase separated droplets of DDX39A (10 µM) within 1 hr incubation at 37°C (**Fig. 6Ci-ii and S6Bi-ii**). Addition of crowding agent PEG resulted in fusion of the droplets (**Fig. S6Bi-ii**). Furthermore, proteostat staining confirmed the amyloidogenic potential of the phase separated protein. Transmission electron microscopy (TEM) also revealed that the size of DDX39A droplets were relatively larger in presence of PEG than in absence of the crowding agent (**Fig. 6Di-vi**). Fusion of smaller droplets was also observed in the PEG-treated samples (**Fig. 6Dvi, arrowhead**). Next, we added H1N1G4.3 rG4 to the phase separated DDX39A. Increased 6-FAM fluorescence suggested that DDX39A retained rG4-helicase activity at the phase separated state, even in presence of PEG when the droplets were fused to each other (**Fig. 6E-F**).

Addition of DDX39A at 1:200 molar ratio moderately enhanced LLPS of 100 µM α-Synuclein suggesting a cooperativity of phase separation between these proteins (**Fig. S5D**). To investigate further, we incubated 10 µM DDX39A tagged with AF488 with increasing concentration of rhodamine-tagged α-Synuclein. Increasing staining of DDX39A droplets with red fluorescence indicated that co-condensation with α-Synuclein (**Fig. 6G**). The droplets were found to be increasingly fused in presence of α-Synuclein, than DDX39A alone (**Fig. 6Ci and iii, 6G**). PEG-mediated fusion of condensates was also more prominent in presence of α-Synuclein (**Fig. 6Cii and iv**). Similarly, numerous small, few larger, and a few fused condensates were observed in TEM images when both the proteins were incubated together (**Fig. 6Dvii-ix**). Condensate sizes were even larger in presence of PEG (**Fig. 6Dx-xii**). We also observed fibril structures in few fields (**Fig. 6Dxii**). The reaction-mixes were further analyzed on SDS-PAGE followed by western blotting with α-Synuclein and DDX39A antibodies, respectively. Monomeric α-Synuclein band was reduced in intensity with concomitant appearance of a high molecular band suggesting increased phase separation in presence of DDX39A and PEG (**Fig. S6C, \***). Upon probing for DDX39A, co-incubation with α-Synuclein resulted in increased intensity of high molecular weight bands in absence of PEG (**Fig. S6C, arrowheads**) while addition of PEG indicated high molecular weight entities throughout (**Fig. S6C, bracketed**). DDX39A helicase activity was not only retained rather enhanced by ∼50% at the phase separated condensates with α-Synuclein as revealed by increased 6-FAM fluorescence (**Fig. 6H**). Interestingly, microscopy indicated formation of fibrillar structures during the assay time point i.e. 16 hr after addition of H1N1G4.3 rG4 in the reaction mixes (**Fig. 6I**). The fibrils were longer in presence of α-Synuclein. We also observed multiple dots along the fibrils co-stained by both proteostat and 6-FAM suggesting hotspots for helicase activity and RNA-mediated initiation of amyloidogenesis (**Fig. 6I**).

Together, our results suggested that DDX39A and α-Synuclein facilitated LLPS of each other; enzymatic activity of DDX39A was elevated during co-phase separation with α-Synuclein; however, amyloidogenesis was concomitantly initiated. Notably, if the tripartite α-Synuclein-viral rG4-DDX39A interaction equilibrium was misbalanced due to excess abundance of viral rG4s, amyloidogenesis was expedited (**Fig. 3I**). While, over-expression of DDX39A in the same cells could counteract the accelerated amyloid formation (**Fig. 4I**).

## Discussion

Several RNA binding proteins (RBPs), such as TDP-43, FUS, and hnRNPA1, contain low complexity domains that facilitate their liquid–liquid phase separation (LLPS) to nucleate pathogenic amyloid fibrils associated with neurodegenerative diseases^26,28,93^. α-Synuclein lacks a canonical RNA-binding domain but it can still associate with RNA and RBPs either directly or through deposition into cytosolic P-bodies^43,44,94^. Accordingly, α-Synuclein-RNA-RBP crosstalk disrupts RNA-protein granule formation and perturbs RNA metabolism. Yet, it remains largely unknown how RBPs that typically regulate RNA secondary structures within these granules and thereby reprogram mRNA transcription might directly alter α-Synuclein amyloidogenesis and subsequent cellular homeostasis.

Here, we report that a DEAD-box RNA-helicase DDX39A, which is a predominantly nuclear protein involved in mRNA splicing and export^74^, localize to cytosolic P-bodies and interact with α-Synuclein in a RNA-dependent manner. We further demonstrate that this interaction decelerates rG4-mediated α-Synuclein amyloid formation by arresting the protein in a granular, amyloid-prone intermediate state. DDX39A itself is intrinsically amyloidogenic and has been identified as a component of RNA-seeded sub-nuclear condensates, known as amyloid bodies (A-bodies)^90^. Our experiments suggest that cooperative phase separation with α-Synuclein promotes helicase activity of DDX39A to unwind rG4 structures. Although DDX39A-mediated unwinding of rG4s transiently limits LLPS of α-Synuclein, we posit that amyloidogenesis is eventually triggered as a long-term trade-off within this tripartite RNA-protein interaction equilibrium (**Fig. 7**). Importantly, DDX39A is highly expressed across multiple brain regions. Mutations in this protein have been implicated in early-onset neurodegenerative diseases associated with cerebral atrophy^95^. Furthermore, our analysis of published mass spectrometry data revealed that DDX39A interacts with soluble α-Synuclein isolated from post-mortem brain sections of Parkinson’s Disease (PD) and other Synucleinopathies^96^ (**Table. S2**).

**Fig. 7:**
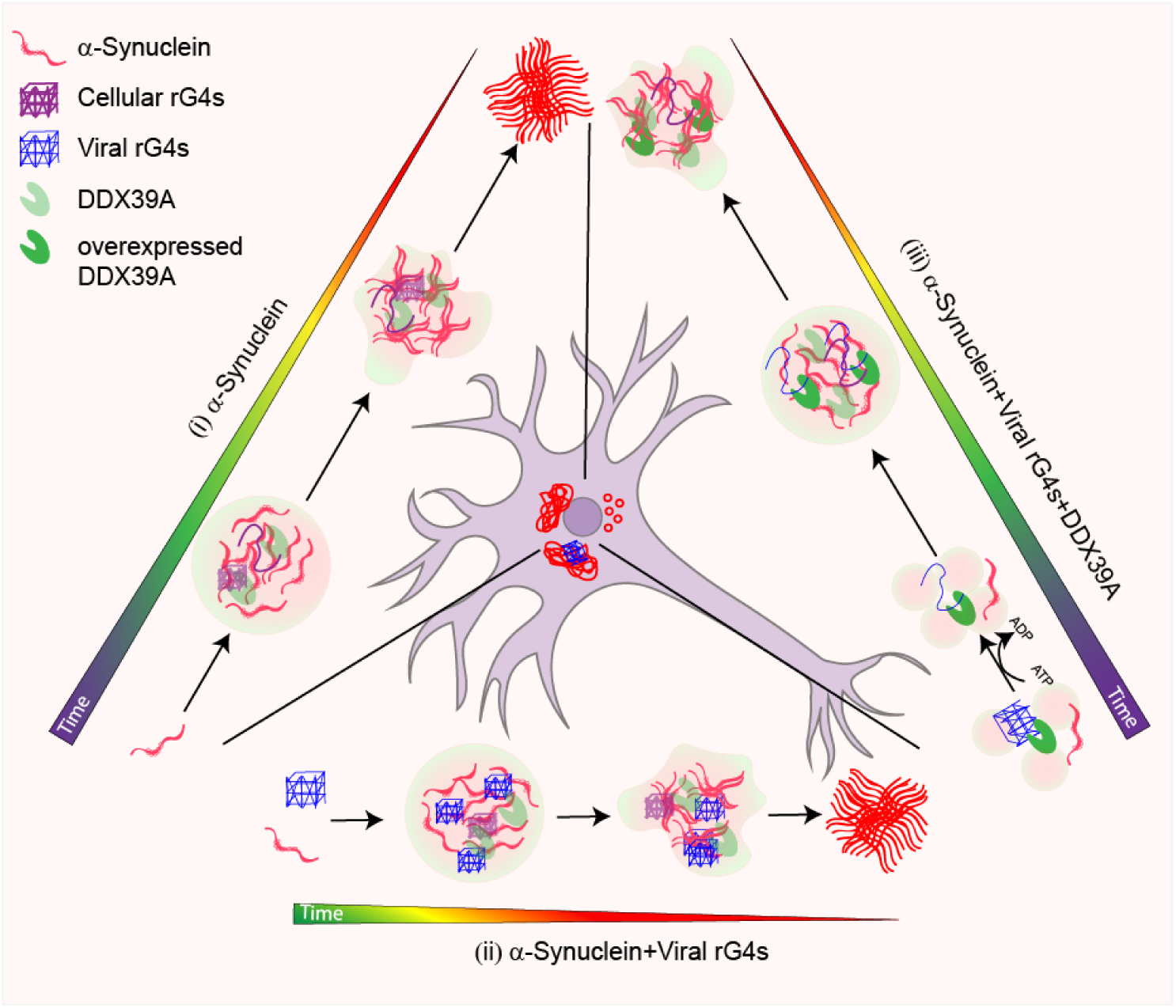
Proposed mechanism of α-Synuclein amyloidogenesis mediated by cytosolic interactions between α-Synuclein, DDX39A, and viral RNA G-quadruplexes (rG4s). (i) α-Synuclein amyloid formation nucleates through liquid-liquid phase separation (LLPS) where condensates mature into more gel like oligomers and finally to filamentous amyloids following a defined timeline due to an equilibrium between cellular rG4s and endogenous DDX39A. (ii) Increased viral rG4s expedites the process of sol-gel transition of α-Synuclein leading to accelerated amyloidogensis. (iii) Overexpression of RNA helicase DDX39A through its ATP dependent rG4-unwinding activity counteracts the expedited amyloidogenesis mediated by viral rG4s. DDX39A helicase activity is enhanced due to its co-condensation with α-Synuclein. However, α-Synuclein continues to form amyloids as a trade-off within this tripartite interaction equilibrium.

DDX39A has also been implicated in the mechanisms of viral infections. Reduced sumoylation of DDX39A in vesicular stomatitis virus (VSV) infected cells impairs nuclear export of mRNAs facilitating VSV-escape from innate immunity^77^. Further, DDX39A binds to secondary structures of viral RNA at the cytoplasm of chikungunya-infected cells to prevent viral propagation^74^. Importantly, viral infections has recently been identified as a long-term risk factor for PD, which is characterized by deposition of α-Synuclein amyloid aggregates in brain^49^. Viral etiology to PD was first proposed with the emergence of a parkinsonian disorder after the 1918 influenza pandemic^97^. Many other case-control studies also proposed that influenza infection might increase the long-term risk of developing PD^50,97^. A mechanistic study in 2020 proposed that disruption of cellular proteostasis by H1N1 influenza may promote α-Synuclein aggregation^98^. Our results, interestingly, offers an alternative mechanism that rG4s from the H1N1 influenza genome can expedite α-Synuclein amyloidogenesis and thereby contribute to PD-pathology. Although influenza virus RNA is largely coated by nucleoprotein (NP), it still forms secondary structures and participates in both intra- and inter-segment RNA–RNA interactions. Genome-wide analyses of RNA structure and NP binding sites show that NP associates with H1N1 viral RNA in a non-uniform manner, creating regions of low NP occupancy where RNA secondary structures are preserved and functionally important for genome organization and packaging^99,100^. We also demonstrate that rG4s from SARS-CoV-2 and JEV genome can potentiate α-Synuclein amyloidogenesis. Indeed, a recent study reports that SARS-CoV-2 infection exacerbates the cellular pathology of PD in human dopaminergic neurons and in a mouse model^79^.

Being abundantly expressed in brain, DDX39A may associate with PD-related α-Synuclein amyloidogenesis by unwinding endogenous mRNA G4s as well as viral rG4s, particularly during JEV infection or in severe cases of influenza and SARS-CoV-2 infection where viral particles are known to reach the brain^98,101^. However, both influenza and SARS-CoV-2 predominantly infect respiratory tracts. In these cases, even if local α-Synuclein amyloidogenesis happens, how it may reach neurons in brain remains an open question. Intriguingly, recent reports suggest that injection of α-Synuclein amyloid fibrils into the gastrointestinal tract and kidney could reach brain inducing LB-related pathology^102–106^. Evidences further suggest that red blood cells might facilitate the inter-organ propagation of α-Synuclein amyloid fibrils^102^. Further, bidirectional propagation of α-Synuclein amyloids from peripheral organs to brain and *vice versa* is reported^107,108^. Thus, initiation of local amyloidosis in peripheral organs by viral infection, or by mRNA vaccines that may contain potential rG4s, and subsequent retrograde transport of α-Synuclein amyloid fibrils to brain cannot be ruled out. In this context, it is also interesting to note that α-Synuclein itself can mitigate RNA virus infections in brain including JEV^109,110^. DDX39A can also alleviate chikungunya infection^74^. Therefore, cooperative phase separation of DDX39A and α-Synuclein in mitigating viral infections is a possibility and will be interesting to investigate using *in vivo* models.

## Methods

### Generation of Expression Constructs

To generate SNCA(DM)-EGFP localization variants (SNCA(DM)-EGFP-NLS, SNCA(DM)-EGFP-NES), localization signals (NLS, NES) were amplified using primers in **Methods Table S1** from **pcDNA/syn^NLS^** and **pcDNA/syn^NES^** plasmids provided by Mel B. Feany lab^111^ as a kind gift and were cloned into pcDNA4/TO-SNCA(DM)-EGFP using BsrGI and XbaI cloning sites. To generate SNCA(DM)-EGFP-KRAS, localization signals (KRAS) from Venus-KRAS-N1^112^ were cloned into pcDNA4/TO-SNCA(DM)-EGFP using BsrGI and XbaI cloning sites. To prepare DDX39A construct, DDX39A sequence was PCR amplified from cDNA prepared from HEK293T cells using primers in **Methods Table S1** and cloned into pCDNA3.1 using KpnI and XbaI. mCherry sequence was amplified from pcDNA3.1-Fluc(DM)-mCherry^13^ and cloned at the C-terminal using Xba1 and Apa1 sites. DDX5 construct was similarly prepared by amplifying the sequence from HEK293T cDNA using primers in **Methods Table S1** and cloned into pCDNA3.1 using KpnI and BamHI sites. mCherry sequence was digested out from pcDNA3.1-Fluc(DM)-mCherry and cloned at the C-terminal with BamHI and XbaI sites. For protein purification, Human SNCA was cloned into pET-28a(+) expression vector using the restriction enzymes HindIII and XhoI and DDX39A was cloned into pET-21b(+) expression vector using HindIII and XhoI^13^.

For lentiviral expression construct, SNCA(WT)-EGFP, SNCA(WT)-EGFP-NES were cloned into pFGE plasmid (gift from Jyotsna Dhawan lab) using AgeI and EcoRI.

### α-Synuclein Protein Purification

α-Synuclein was purified based on previously published protocol^13^. Briefly, SNCA-pET-28a(+) construct were transformed into *E. coli* BL21(DE3) and induced by adding 1 mM IPTG (Sigma) at 0.6 OD_600_. The cells were pelleted 3 hr after induction, then resuspended in TNE buffer (50 mM Tris, 150 mM NaCl, 10 mM EDTA pH 8), and frozen in – 80°C until use. Frozen cell suspensions were subjected to heat shock at 95°C for 10 min and centrifuged at 18,000 g. From the supernatant, DNA was precipitated using 10% (w/v) streptomycin sulfate and glacial acetic acid followed by protein precipitation from the supernatant by saturated ammonium sulfate. Thereafter, the pelleted protein was washed thrice by 100 mM ammonium acetate and further precipitated using 100% ethanol. The precipitate was dissolved in HEPES buffer (50 mM HEPES, 100 mM KCl pH 7) and size exclusion chromatography was performed using HiLoad 16/600 Superdex 75 pg Column (GE28-9893-33, GE Healthcare) attached to BioLogic DuoFlow Chromatography system (Bio-Rad). Purified α-Synuclein was concentrated using 10 kDa Amicon (Merck Millipore) up to 1 mg/ml in HEPES buffer for preformed fibril (PFF) generation or storage at – 80°C.

### Protein labelling of α-Synuclein

Rhodamine labelling was done as per manufacturer’s instructions (ThermoFisher Scientific, USA). In brief, 10X molar excess of NHS-Rhodamine (dissolved in DMSO) was added to the protein. The mixture was incubated at 4°C for 4 hours with continuous slow rotation. The excess dye was removed using SpinOut desalting columns. 1:19 molar ratio of labeled versus unlabeled protein was used for experiments.

### Preformed fibril (PFF) generation

1 ml of concentrated protein (1 mg/ml) was aliquoted into 1.5 ml tubes (LoBind, Eppendorf) and kept on Eppendorf ThermoMixer C at 37°C with 1500 rpm for 48 hr. Fibril formation was checked using a small aliquot in Thioflavin T (ThT) assay^13^. The formed fibrils were sonicated using 3 mm thick probe sonicator (cat: VCX 130 PB, SONICS Vibra-Cell) for 1 min with setting: 5 second ON, 2 second OFF, 15 % amplitude. Sonicated PFF were then aliquoted into 50 and 100 µl and stored in – 80°C until use. The PFF stored at – 80°C were stable up to 5-6 months with similar aggregation propensity in cell culture.

### DDX39A Protein Purification

DDX39A-pET-28a(+) construct were transformed into *E. coli* Rosetta(DE3) and induced by adding 1 mM IPTG (Sigma) at 0.7 OD_600_. The cells were pelleted 16 hr after induction at 18°C and was resuspended in lysis buffer (50 mM Tris, 250 mM KCl, 10 mM 2-mercaptoethanol, pH 8.0). Cells were lysed by sonication using probe sonicator (HBSNII92 650W Ultrasonicprobe sonicator) for 30 min with setting: 20 second ON, 20 second OFF, 30 % amplitude. The lysate was cleared by centrifugation at 12,000×g for 10 minutes. Lysate was applied to a packed Ni-NTA agarose bead column equilibrated with lysis buffer (50 mM Tris, 250 mM NaCl, 10 mM 2-mercaptoethanol, pH 8.0). The column was washed sequentially with wash buffers: 1) 50 mM Tris/HCl, 1 M KCl, 10 mM 2-mercaptoethanol, pH 8. 2) 50 mM Tris/HCl, 1 M KCl, 10 mM 2-mercaptoethanol, 10 mM Imidazole, pH 8. Elution was with 50 mM Tris/HCl, 1 M KCl, 10 mM β-Me, 250 mM Imidazole, pH 8.0.

### Protein labelling of DDX39A

Alexa fluor 488 labelling was done as per manufacturer’s instructions (ThermoFisher Scientific, USA). In brief, 10X molar excess of Alexa flour 488 C5-maleimide (dissolved in DMSO) was added to the protein. The mixture was incubated at 4°C overnight with continuous slow rotation. The excess dye was removed using SpinOut desalting columns. 1:9 molar ratio of labeled versus unlabeled protein was used for experiments.

### Cell Culture, transfection and preparation for microscopy

HEK293T cells were maintained in Dulbecco’s Modified Eagle’s Medium (Gibco) supplemented with 10 % fetal bovine serum (FBS) (Gibco) and 90 U/ml penicillin (Sigma) 50 µg/ml streptomycin (Sigma) at 37°C and 5 % CO_2_. Transfection of cells was performed with Lipofectamine 3000 reagent (Invitrogen) according to the manufacturer’s protocol. For stable cell line preparation, pcDNA6/TR and pcDNA4/TO system was used (Life Technologies). One day after transfection with pcDNA6/TR, 10,000 cells were seeded in a 100 mm dish and selected with 5 µg/ml Blasticidin S HCl (Invitrogen). Colonies formed after 2-3 weeks were analyzed for anti-TET repressor by western blots (data not shown). Positive clone for pcDNA6/TR was transfected with various pcDNA4/TO constructs (SNCA(DM)-EGFP, SNCA(DM)-EGFP-NLS, SNCA(DM)-EGFP-NES and SNCA(DM)-EGFP-KRAS) and selected with 200 µg/ml Zeocin (Invitrogen). Surviving colonies after 2-3 weeks were tested for induction by adding 1 µg/ml Doxycycline (MP Bioscience) by fluorescence microscopy and western blot. Positive clones were frozen and/or maintained for further experiment.

Induction of α-Synuclein aggregation – stable cell lines for SNCA(DM)-EGFP, SNCA(DM)-EGFP-NLS, SNCA(DM)-EGFP-NES and SNCA(DM)-EGFP-KRAS were induced by 1 µg/ml Doxycycline for 24 hr. Next day, induced cells were trypsinized and subcultured into various dishes according to **Method Table S2** for 6 days. 1 µM PFF and 1 µg/ml doxycycline was added after 6 hr of cell plating.

RNA treatment to cell culture – stable cell lines for SNCA(DM)-EGFP, SNCA(DM)-EGFP-NLS, and SNCA(DM)-EGFP-NES were plated into various dishes according to **Method Table S2**. Next day, cells were transfected with RNA using Lipofectamine RNAimax reagent (Invitrogen). 1 µM PFF and 1 µg/ml doxycycline was added after 24 hr and cell were cultured for 3 days.

DDX39A overexpression experiments – pcDNA3.1 constructs with mCherry, DDX39A-mCherry and DDX5-mCherry were transfected into the stable lines as indicated in results section using Lipofectamine3000 reagent (Invitrogen). 1 µM PFF and 1 µg/ml doxycycline was added after 24 hrs and cell were cultured for 6 days.

Acetone/Methanol fixation – Cells grown on cover glass were washed with PBS to remove medium. Pre-chilled acetone and methanol in ratio 1:1 was added to the cells and plate was incubated at -20 °C for 5 min followed by PBS wash. Paraformaldehyde fixation – Cells were incubated with 4% paraformaldehyde solution at room temperature for 15 min followed by PBS wash. 0.1% Triton X100 in PBS was used for cell permeabilization prior to immunostaining.

### Primary Neuron Culture

Hippocampal Primary neurons were isolated from E17-E19 C57BL/6 mouse brains (Charles River Laboratories) as described in Seibenhener et al.^113^. For microscopy, approx. 120,000 cells were cultured on poly-D-lysine coated 18 mm glass coverslips and 300,000 cells were cultured on 6-well plates for other cell biology experiments. Neurobasal Feeding Media was replaced every third day.

Induction of α-Synuclein aggregation – 100 nM PFF was added to neuron cultures at 5 days *in vitro* (DIV) and incubated for 10 days at described in Volpicelli-Daley et al.^114^. RNA treatment – neurons were transfected using Lipofectamine2000 reagent (Invitrogen) at 5 DIV. 100 nM PFF was added after 24 hrs and neurons were cultured for another 4 DIV. Co-transfection of RNA and DDX39A – neurons were transfected with pcDNA3.1 constructs (mCherry, DDX39A-mCherry and DDX39AΔRNA-mCherry) and RNA using Lipofectamine2000 reagent (Invitrogen) at 5 DIV. 100 nM PFF was added after 24 hr and cell were cultured for another 4 DIV. Neurons were fixed for microscopy with 1:1 acetone and methanol chilled at -20°C and immunostained with antibodies.

All experiments were performed in accordance with the standard protocols and procedures approved by the Institutional Animal Ethics Committee (IAEC).

### Lentivirus production and transduction in neurons

For viral production, HEK293T cells were transfected by using Lipofectamine3000 reagent (Invitrogen) with the expression (pFGE) vector and two helper vectors, psPAX2, pHCMV-EcoEnv at 24, 12, and 8μg of DNA respectively per 15-cm plate. The supernatants of plates were collected every day after 24hr to 72hrs and pooled, spun at 500 × g for 15 min. Clarified supernatant was concentrated using Lenti-X concentrator (Takara Cat. Nos. 631231) according to product specific protocols. In brief, Lenti-X concentrator was added to the clarified supernatant in 1:3 ratio and incubated for 48 hours, spun at 1500×g for 45 minutes. The pellet was resuspended in 1/100 times PBS, aliquoted and stored at – 80°C until use. Neurons were transduced at DIV7 by thawing the viral particles in serum free Neurobasal media followed by to the cultures using the previously described protocol^115^.

### Viral RNA isolation

MDCK cells were infected at a multiplicity of infection (MOI) of 0.01 with influenza A virus A/Human/WSN/1933 (H1N1), and cytopathic effect (CPE) was monitored to assess infection progression. At 48 hours post-infection, when approximately 70–80% CPE was observed, cell culture supernatants containing high titers of virus (∼2 × 10⁸ plaque-forming units [PFU]/mL) were harvested and clarified by sequential centrifugation at 5,000 × g for 15 min followed by 20,000 × g for 20 min to remove cell debris. Clarified supernatants were filtered through a 0.22-μm membrane and subjected to ultracentrifugation at 150,000 × g for 2 h at 4 °C in sterile Quick-Seal round-top polypropylene tubes using a Type 70 Ti fixed-angle rotor (Beckman Coulter) ref#. The resulting virus pellet was directly resuspended in TRIzol™ reagent (Invitrogen) for RNA isolation according to the manufacturer’s instructions. Purified RNA was resuspended in nuclease-free water, and RNA yield was assessed spectrophotometrically^116^.

### Viral Infection and Titration

Mice primary hippocampal neurons were infected with the SARS-CoV-2 B.1.1.8 variant at a multiplicity of infection (MOI) of 0.1 in serum-free feeding medium, 24 h post-transfection. Following 72 h of infection, culture supernatants were collected for viral titration by plaque assay. Cells were subsequently harvested and fixed for immunoblotting and confocal microscopy analyses.

### Plaque Forming Assay

Viral titers in the collected supernatants were determined by plaque forming assay on Vero cells. Supernatants were serially diluted in serum-free medium (SFM) and used to infect Vero cell monolayers. After 3 h of viral adsorption, the inoculum was removed and cells were overlaid with a 1:1 mixture of 2× DMEM and 1% low-melting agarose (LMA) prepared in water. Plates were incubated undisturbed for 6–7 days. Cells were then fixed with 4% formaldehyde, the agarose overlay was removed, and plaques were visualized by staining with 0.1% crystal violet and the dilution which had 5–20 plaques were used for calculating PFU/mL.

### Immunofluorescence microscopy

Cells were grown on 18 mm glass coverslips in 12 well plate (Nunc) and fixed as explained above. After fixation, blocking was done by incubating the coverslips with 5% BSA prepared in PBS with 0.05% tween-20 (PBST) for 30 min at RT on shaker. Required primary antibody (**Methods Table S3**) was added and coverslips were kept at 4°C overnight followed by PBST wash. Suitable secondary antibody was added and incubated for 1 hr at room temperature. Antibody dilutions were prepared in 0.1% BSA with PBST as mentioned in **Methods Table S3.** Slides were prepared after DAPI staining using 10 µl Antifade Mounting Medium (cat: H-1000, Vectashield). Imaging was done in Leica TCS SP8 laser scanning microscope.

### Cell lysis, SDS-PAGE and Western Blotting

Cell pellets were lysed in TBS-T lysis buffer (1% Triton X-100 in 50 mM Tris-HCl, 150 mM KCl, and a cocktail of protease inhibitors (Roche)) at 4°C and incubated for 1 hr with intermittent vortexing. Lysed cells were centrifuged at 12,000 g for 15 min at 4°C, and the supernatant was collected as soluble fraction. The pellet was washed twice with 1×PBS and boiled in 4×SDS loading buffer (0.2 M Tris–HCl pH 6.8, 8 % SDS, 0.05 M EDTA, 4 % 2-mercaptoethanol, 40 % glycerol, 0.8 % bromophenol blue) for 15 min to obtain the insoluble fraction. Total fraction was prepared by directly boiling the cell pellet in 4×SDS loading buffer for 15 min. Protein estimation was carried out by Amido Black Protein assay. Protein fractions were separated by SDS-PAGE and transferred onto 0.2 μm PVDF membrane (Bio-Rad) for 100 min at 300 mA using the Mini-Trans Blot cell system (Bio-Rad). Membranes were probed by primary and secondary antibodies (**Methods Table S3**) and imaged using Chemidoc MP imaging system.

For Primary neurons, total fraction was collected by directly lysing the cells with 4×SDS loading buffer (0.2 M Tris–HCl pH 6.8, 8 % SDS, 0.05 M EDTA, 4 % 2-mercaptoethanol, 40 % glycerol, 0.8 % bromophenol blue) and boiled for 15 minutes. Protein estimation was carried out by Amido Black Protein assay. Protein fractions were separated by SDS-PAGE and transferred onto 0.2 μm PVDF membrane (Bio-Rad) for 100 min at 300 mA using the Mini-Trans Blot cell system (Bio-Rad). The membrane was fixed with 4% Paraformaldehyde solution for 45 minutes and washed with PBS before blocking. Membranes were probed by primary and secondary antibodies (**Methods Table S3**) and imaged using Chemidoc MP imaging system.

### Immunoprecipitation and sample preparation for Mass Spectrometry

Cell pellets were lysed in TBS-T lysis buffer (1% Triton X-100 in 50 mM Tris-HCl, 150 mM KCl, and a cocktail of protease inhibitors (Roche)) at 4°C and incubated for 1 hr with intermittent vortexing. Lysed cells were centrifuged at 12,000 g for 15 min at 4°C, and the supernatant was collected as soluble fraction. Immunuprecipitation was done using µMACS™ GFP Isolation Kit (Cat- 130-091-125). Equal amount of soluble fraction was incubated with µMACS anti-GFP MicroBeads for 4 hrs. Labelled cell lysate was applied to the MACS column and washed according to the manufacturer’s protocol. Target proteins were eluted by adding 4×SDS loading buffer (0.2 M Tris–HCl pH 6.8, 8 % SDS, 0.05 M EDTA, 4 % 2-mercaptoethanol, 40 % glycerol, 0.8 % bromophenol blue) preheated at 95°C.

Eluted fractions were separated on NuPAGE 4%–12% Bis–Tris Protein Gels (Invitrogen) in MES buffer (100 mM MES, 100 mM Tris–HCl, 2 mM EDTA, 7 mM SDS) at 200 V for 40 min, fixed and stained with Coomassie brilliant blue. Preparation of gel slices, reduction, alkylation, and in-gel protein digestion was carried out as described earlier^117,118^. Finally, peptides were desalted and enriched according to Rappsilber et al.^119^ and stored in - 20°C until mass spectrometry analysis.

### Analysis of mass spectrometry data

For label-free quantification, raw spectra files were loaded onto the MaxQuant platform (Ver. 1.6.10.43) and searched against the SwissProt *Homo sapiens* database (2024) with Label Free Quantification (LFQ) option. Search parameters included static and dynamic modification of cysteine by carbamidomethylation and dynamic modification of methionine by oxidation. Enzyme specificity of trypsin allowing for up to two missed cleavages. Precursor mass tolerance was set to 5 ppm and fragment mass tolerance to 0.02 Da. Other parameters included a minimum of one peptide for identification with six amino acid length. Percolator (q-value) was used for validating peptide spectrum matches accepting only the top-scoring hit for each spectrum, satisfying the cut-off values for false discovery rate of 1%. Experiments were performed in triplicate. Proteins quantified in at least two replicates were included in the analysis. We considered the identified proteins to be α-Synuclein–interacting proteins if they met the following conditions: the peptides were detected in at least two of three repeat experiments with SNCA(DM)-EGFP and/or its localization variants but not in corresponding EGFP-controls. For the proteins quantified in at least two repeats of both SNCA(DM)-EGFP and EGFP-controls, Log2 fold change was calculated from the ratios of LFQ intensities of between the experiment and control from average LFQ intensities of three independent experiments. Graphs were prepared in Perseus. GraphPad Prism v. 9.4.1. GO analysis of the quantified proteins was performed using SHINY GO 0.80 version^120^. Venn Diagrams were prepared using https://bioinformatics.psb.ugent.be/webtools/Venn/ web-tool. RNA binding proteins and mRNA binding proteins were mapped using RBPGO^121^.

### Circular Dichroism spectroscopy

RNA oligonucleotides were synthesized commercially by GenScript (USA). Each RNA (3μM) was prepared in 10 mM Tris-HCl,150 mM KCl buffer (pH 7.4). The oligomers were then refolded by a heating/cooling process (95°C for 5 min, followed by gradual cooling to room temperature overnight) before measurement. The circular dichroism spectra were recorded at room temperature over the range of 220–320 nm using a spectrometer (JASCO Corp., J-810, Rev. 1.00) equipped with a 3-mm path-length quartz.

### LLPS of α-Synuclein

100 μM α-Synuclein (95% unlabelled and 5% labelled) was incubated with 1 μM RNA and 0.5 μM DDX39A, either with or without 5 mM ATP and 5 mM MgCl_2_. Reactions were set up in HEPES- KCl Azide buffer (50 mM HEPES, 150 mM KCl, 0.04% sodium azide, pH 8). PEG 8000 was added to all reaction mixtures at a final concentration of 20% (v/v) to induce molecular crowding. The reactions then incubated at 37°C. At time points of 30 minutes, 2 hours, and 24 hours, aliquots were taken from each reaction mixture and followed by imaging using confocal microscopy and Transmission electron microscopy. For all *in vitro* experiments, synthesized RNA oligo was used and both the proteins were purified in the lab using bacterial overexpression system.

### Confocal Microscopy and FRAP of LLPS

5 μl of the LLPS reaction mix was drop-casted on acid washed glass slides and mounted using 18 mm round coverslips. For proteostat staining, Proteostat Dye (catalog no. ENZ-51035) was diluted 20 times and added to the sample in 1:5 ratio. For FRAP assay, samples were photobleached with 100% laser power, and time-lapse images were recorded every 0.7s using the LSM880 microscopy system (Carl Zeiss) with a 63× oil objective (NA 1.4) and a 568-nm laser pulse on the region of interest. Imaging was done in Leica TCS SP8 and Ziess LSM880 laser scanning microscope.

### Transmission electron microscopy

LLPS reaction mix was diluted 100 times and 3.5 μl of sample was spotted onto a 300 mesh copper grid (2005C-XA, SPI supplies) and incubated at room temperature (RT) for 1 min. Thereafter, the remaining sample was gently wiped off with Whatman® filter paper. This was followed by staining of all the samples with 3.5 µl of uranyLess (catalog no. - 22409) solution (2% w/v) for another 1 min and the remaining sample was gently wiped off with Whatman® filter paper. Finally, the grids were air-dried for 15 min. Images were acquired using a 120 kV transmission electron microscope (Thermo Scientific Talos L120C) accelerated at 120 kV.

### Electrophoretic Mobility Shift Assay

To study the binding of RNA and DDX39A, RNA stock was diluted in buffer (10 mM Tris-Cl, 150 mM KCl, pH 7.4) and denatured by heating at 95°C for 5 minutes, followed by gradual cooling to room temperature to promote proper folding. For setting up the reactions, 500 nM RNA was incubated with increasing concentrations of DDX39A (100 nM, 250 nM, 500 nM) either in presence or absence of 5 mM ATP and 5 mM MgCl_2_ in reaction buffer (50 mM Tris-Cl, 250 mM KCl, 10 mM 2-mercaptaethanol, pH 8.0). Reactions were incubated for 4 hours at RT and 37°C in parallel. Following incubation, samples were mixed with native RNA loading dye and resolved on 8% native polyacrylamide gel in 0.5X TBE buffer. The gel was subsequently stained with SYBR Gold to visualize RNA bands and RNA-protein complexes.

### DDX39A Helicase Activity

250 nM of recombinant purified DDX39A or 250 nM BSA were incubated with 500 nM RNA for 16 hours at 37°C either in presence or absence of 5 mM ATP and 5 mM MgCl_2_ in reaction buffer (50 mM Tris-Cl, 250 mM KCl, 10 mM 2-mercaptoethanol, pH 8.0). The reaction was setup using RNA that had been preheated at 95°C for 5 minutes and then gradually cooled to room temperature. After 16 hours of incubation, reaction mixtures were mixed with native RNA loading dye and resolved on 10% native polyacrylamide gel in 0.5X TBE buffer. The gel was subsequently stained with SYBR Gold to visualize RNA bands.

To assess the helicase activity of DDX39A using Fluorescence resonance energy transfer (FRET), 20 nM of recombinant purified DDX39A were incubated with 40 nM RNA (conjugated with a FAM signal at 5’ end and BHQ group at 3’ end) for 16 hours at 37°C either in the presence or absence of 350 μM ATP and 350 μM MgCl_2_ in reaction buffer (50 mM Tris-Cl, 250 mM KCl, 10 mM 2-mercaptaethanol, pH 8.0). The reaction was setup using RNA that had been preheated at 95°C for 5 minutes and then gradually cooled to room temperature. After 16 hours of incubation, fluorescence changes were monitored using an excitation wavelength of 490 nM and emission at 520 nM.

### DDX39A Phase Separation

To investigate the liquid-liquid phase separation of DDX39A, protein was labelled with Alexa flour 488 C5-maleimide. Reaction was set with increasing concentration of DDX39A (90% unlabelled and 10% labelled) as indicated in **Fig. 6** either in presence or absence of PEG 8000. Reactions were set up in reaction buffer (50 mM Tris-Cl, 250 mM KCl, 10 mM 2-mercaptaethanol, pH 8.0) and PEG 8000 was added at a final concentration of 10% (v/v). The reactions then incubated at 37°C. After 1 hour of incubation, aliquots were taken from each reaction mixture and followed by imaging using confocal microscopy.

To investigate LLPS of DDX39A in the presence of α-Synuclein, both the proteins were labelled with Alexa flour 488 C5-maleimide and NHS-Rhodamine, respectively. 10 μM DDX39A (90% unlabelled and 10% labelled) was incubated with increasing concentration of α-Synuclein as indicated in **Fig. 6** either in presence or absence of PEG. Reactions were set up in reaction buffer (50 mM Tris-Cl, 250 mM KCl, 10 mM 2-mercaptaethanol, pH 8.0) and PEG 8000 was added at a final concentration of 10% (v/v). The reactions then incubated at 37°C. After 1 hour of incubation, aliquots were taken from each reaction mixture and followed by imaging using confocal microscopy.

### DDX39A Helicase Activity after Phase Separation

To assess the helicase activity of DDX39A after phase separation using FRET, liquid-liquid phase separation of 10 μM DDX39A was incubated with or without 10 μM α-Synuclein either in presence or absence of PEG 8000 for one hour. 0.6 μl of these reaction mixtures were incubated with 40 nM RNA (conjugated with a FAM signal at 5’ end and BHQ group at 3’ end) for 16 hours at 37°C in the presence of 150 μM ATP and 150 μM MgCl₂ in reaction buffer (50 mM Tris-Cl, 250 mM KCl, 10 mM 2-mercaptaethanol, pH 8.0). Final concentration of DDX39A in the helicase reaction mixture was maintained at 20 nM.

### Image processing and quantification

To count perinuclear amyloid positive cells, z-stack images were obtained using Leica SP8 confocal microscope with a 63× oil objective (NA 1.4). Positive cells within the obtained images were manually counted.

Quantification of DCP1A-positive P-bodies was performed using Fiji/ImageJ (v1.53 or later) using a custom macro. Individual cell boundaries were manually segmented and added as single ROIs to the ROI Manager. A pre-generated binary mask of DCP1A-positive P-bodies (white objects on a black background) was used as the input image. For each cell boundary ROI, Analyze Particles was applied with a size threshold of 0.05–2 μm^2^ to quantify the number of DCP1A-positive P-bodies and to extract individual P-body area measurements. For each cell, the total P-body area was calculated as the sum of all detected particle areas, and the average P-body size was calculated by dividing total area by particle number. Aggregate statistics, including total P-body counts, cumulative area, and overall average P-body size, were compiled across all nuclei. All measurements were exported to the ImageJ Log window for downstream analysis.

## Notes

### Competing Interest Statement

The authors have declared no competing interest.

## References

1. Curran, S. & Wrigley, M. Lewy bodies. Am J Psychiatry 154, 1322–3 (1997).

2. Saudou, F. & Humbert, S. The Biology of Huntingtin. Neuron 89, 910–26 (2016).

3. Giasson, B.I. et al. Initiation and synergistic fibrillization of tau and alpha-synuclein. Science 300, 636–40 (2003).

4. Treusch, S., Cyr, D.M. & Lindquist, S. Amyloid deposits: protection against toxic protein species? Cell Cycle 8, 1668–74 (2009).

5. Chiti, F. & Dobson, C.M. Protein misfolding, functional amyloid, and human disease. Annu Rev Biochem 75, 333–66 (2006).

6. Chiti, F. & Dobson, C.M. Protein Misfolding, Amyloid Formation, and Human Disease: A Summary of Progress Over the Last Decade. Annu Rev Biochem 86, 27–68 (2017).

7. Monsellier, E., Ramazzotti, M., Taddei, N. & Chiti, F. Aggregation propensity of the human proteome. PLoS Comput Biol 4, e1000199 (2008).

8. Hipp, M.S., Park, S.H. & Hartl, F.U. Proteostasis impairment in protein-misfolding and - aggregation diseases. Trends Cell Biol 24, 506–14 (2014).

9. Schaffar, G. et al. Cellular toxicity of polyglutamine expansion proteins: mechanism of transcription factor deactivation. Mol Cell 15, 95–105 (2004).

10. Schneider, M.M. et al. The Hsc70 disaggregation machinery removes monomer units directly from alpha-synuclein fibril ends. Nat Commun 12, 5999 (2021).

11. Vabulas, R.M., Raychaudhuri, S., Hayer-Hartl, M. & Hartl, F.U. Protein folding in the cytoplasm and the heat shock response. Cold Spring Harb Perspect Biol 2, a004390 (2010).

12. Landles, C., et al. Subcellular Localization And Formation Of Huntingtin Aggregates Correlates With Symptom Onset And Progression In A Huntington’S Disease Model. Brain Commun 2, fcaa066 (2020).

13. Mansuri, S. et al. Widespread Nuclear Lamina injuries defeat proteostatic purposes of alpha-Synuclein amyloid Inclusions. J Cell Sci (2024).

14. Raychaudhuri, S., Sinha, M., Mukhopadhyay, D. & Bhattacharyya, N.P. HYPK, a Huntingtin interacting protein, reduces aggregates and apoptosis induced by N-terminal Huntingtin with 40 glutamines in Neuro2a cells and exhibits chaperone-like activity. Hum Mol Genet 17, 240–55 (2008).

15. Liu, K.Y. et al. Disruption of the nuclear membrane by perinuclear inclusions of mutant huntingtin causes cell-cycle re-entry and striatal cell death in mouse and cell models of Huntington’s disease. Hum Mol Genet 24, 1602–16 (2015).

16. Hackam, A.S., Singaraja, R., Zhang, T., Gan, L. & Hayden, M.R. In vitro evidence for both the nucleus and cytoplasm as subcellular sites of pathogenesis in Huntington’s disease. Hum Mol Genet 8, 25–33 (1999).

17. Saudou, F., Finkbeiner, S., Devys, D. & Greenberg, M.E. Huntingtin acts in the nucleus to induce apoptosis but death does not correlate with the formation of intranuclear inclusions. Cell 95, 55–66 (1998).

18. Woerner, A.C. et al. Cytoplasmic protein aggregates interfere with nucleocytoplasmic transport of protein and RNA. Science 351, 173–6 (2016).

19. Riguet, N. et al. Nuclear and cytoplasmic huntingtin inclusions exhibit distinct biochemical composition, interactome and ultrastructural properties. Nat Commun 12, 6579 (2021).

20. Yadav, M. et al. Huntingtin is an RNA binding protein and participates in NEAT1-mediated paraspeckles. Sci Adv 10, eado5264 (2024).

21. Morelli, C. et al. RNA modulates hnRNPA1A amyloid formation mediated by biomolecular condensates. Nat Chem 16, 1052–1061 (2024).

22. Hennig, S. et al. Prion-like domains in RNA binding proteins are essential for building subnuclear paraspeckles. J Cell Biol 210, 529–39 (2015).

23. Yan, X. et al. Intra-condensate demixing of TDP-43 inside stress granules generates pathological aggregates. Cell 188, 4123–4140 e18 (2025).

24. Rossi, S. et al. Cytoplasmic accumulation of a splice variant of hnRNPA2/B1 contributes to FUS-associated toxicity in a mouse model of ALS. Cell Death Dis 16, 219 (2025).

25. Oldani, E.G. et al. The effect of G-quadruplexes on TDP43 condensation, distribution, and toxicity. Structure (2025).

26. Molliex, A. et al. Phase separation by low complexity domains promotes stress granule assembly and drives pathological fibrillization. Cell 163, 123–33 (2015).

27. Lester, E. et al. Tau aggregates are RNA-protein assemblies that mislocalize multiple nuclear speckle components. Neuron 109, 1675–1691 e9 (2021).

28. Maharana, S. et al. RNA buffers the phase separation behavior of prion-like RNA binding proteins. Science 360, 918–921 (2018).

29. Hallegger, M. et al. TDP-43 condensation properties specify its RNA-binding and regulatory repertoire. Cell 184, 4680–4696 e22 (2021).

30. Maroteaux, L., Campanelli, J.T. & Scheller, R.H. Synuclein: a neuron-specific protein localized to the nucleus and presynaptic nerve terminal. J Neurosci 8, 2804–15 (1988).

31. Li, W.W. et al. Localization of alpha-synuclein to mitochondria within midbrain of mice. Neuroreport 18, 1543–6 (2007).

32. Hoozemans, J.J. et al. Activation of the unfolded protein response in Parkinson’s disease. Biochem Biophys Res Commun 354, 707–11 (2007).

33. Gosavi, N., Lee, H.J., Lee, J.S., Patel, S. & Lee, S.J. Golgi fragmentation occurs in the cells with prefibrillar alpha-synuclein aggregates and precedes the formation of fibrillar inclusion. J Biol Chem 277, 48984–92 (2002).

34. Lee, H.J., Khoshaghideh, F., Patel, S. & Lee, S.J. Clearance of alpha-synuclein oligomeric intermediates via the lysosomal degradation pathway. J Neurosci 24, 1888–96 (2004).

35. Kowalski, A. et al. Monomeric alpha-synuclein activates the plasma membrane calcium pump. EMBO J 42, e111122 (2023).

36. Goedert, M., Spillantini, M.G., Del Tredici, K. & Braak, H. 100 years of Lewy pathology. Nat Rev Neurol 9, 13–24 (2013).

37. Shults, C.W. Lewy bodies. Proc Natl Acad Sci U S A 103, 1661–8 (2006).

38. Ray, S. et al. alpha-Synuclein aggregation nucleates through liquid-liquid phase separation. Nat Chem 12, 705–716 (2020).

39. Hou, K. et al. Liquid-liquid phase separation regulates alpha-synuclein aggregate and mitophagy in Parkinson’s disease. Front Neurosci 17, 1250532 (2023).

40. Alafuzoff, I. et al. Staging/typing of Lewy body related alpha-synuclein pathology: a study of the BrainNet Europe Consortium. Acta Neuropathol 117, 635–52 (2009).

41. Mahul-Mellier, A.L. et al. The process of Lewy body formation, rather than simply alpha-synuclein fibrillization, is one of the major drivers of neurodegeneration. Proc Natl Acad Sci U S A 117, 4971–4982 (2020).

42. Millett, M. et al. Pathological alpha-synuclein perturbs nuclear integrity. Neurobiol Dis 214, 107028 (2025).

43. Hallacli, E. et al. The Parkinson’s disease protein alpha-synuclein is a modulator of processing bodies and mRNA stability. Cell 185, 2035–2056 e33 (2022).

44. Matsuo, K. et al. RNA G-quadruplexes form scaffolds that promote neuropathological alpha-synuclein aggregation. Cell 187, 6835–6848 e20 (2024).

45. Wang, J. et al. Protein-protein interactions regulating alpha-synuclein pathology. Trends Neurosci 47, 209–226 (2024).

46. Dhakal, S. et al. alpha-Synuclein emulsifies TDP-43 prion-like domain-RNA liquid droplets to promote heterotypic amyloid fibrils. Commun Biol 6, 1227 (2023).

47. Dhakal, S. et al. Distinct neurotoxic TDP-43 fibril polymorphs are generated by heterotypic interactions with alpha-Synuclein. J Biol Chem 298, 102498 (2022).

48. Gracia, P. et al. Molecular mechanism for the synchronized electrostatic coacervation and co-aggregation of alpha-synuclein and tau. Nat Commun 13, 4586 (2022).

49. Levine, K.S. et al. Virus exposure and neurodegenerative disease risk across national biobanks. Neuron 111, 1086–1093 e2 (2023).

50. Cocoros, N.M. et al. Long-term Risk of Parkinson Disease Following Influenza and Other Infections. JAMA Neurol 78, 1461–1470 (2021).

51. Yamada, T. Viral etiology of Parkinson’s disease: Focus on influenza A virus. Parkinsonism Relat Disord 2, 113–21 (1996).

52. Toovey, S., Jick, S.S. & Meier, C.R. Parkinson’s disease or Parkinson symptoms following seasonal influenza. Influenza Other Respir Viruses 5, 328–33 (2011).

53. Conway, K.A. et al. Accelerated oligomerization by Parkinson’s disease linked alpha-synuclein mutants. Ann N Y Acad Sci 920, 42–5 (2000).

54. Conway, K.A. et al. Acceleration of oligomerization, not fibrillization, is a shared property of both alpha-synuclein mutations linked to early-onset Parkinson’s disease: implications for pathogenesis and therapy. Proc Natl Acad Sci U S A 97, 571–6 (2000).

55. Welman, A., Burger, M.M. & Hagmann, J. Structure and function of the C-terminal hypervariable region of K-Ras4B in plasma membrane targetting and transformation. Oncogene 19, 4582–91 (2000).

56. Lu, J. et al. Types of nuclear localization signals and mechanisms of protein import into the nucleus. Cell Commun Signal 19, 60 (2021).

57. Fukuda, M., Gotoh, I., Gotoh, Y. & Nishida, E. Cytoplasmic localization of mitogen-activated protein kinase kinase directed by its NH2-terminal, leucine-rich short amino acid sequence, which acts as a nuclear export signal. J Biol Chem 271, 20024–8 (1996).

58. Laor, D. et al. Fibril formation and therapeutic targeting of amyloid-like structures in a yeast model of adenine accumulation. Nat Commun 10, 62 (2019).

59. Kim, Y.E. et al. Soluble Oligomers of PolyQ-Expanded Huntingtin Target a Multiplicity of Key Cellular Factors. Mol Cell 63, 951–64 (2016).

60. Rostam, N. et al. CD-CODE: crowdsourcing condensate database and encyclopedia. Nat Methods 20, 673–676 (2023).

61. Sanchez de Groot, N., et al. RNA structure drives interaction with proteins. Nat Commun 10, 3246 (2019).

62. Standart, N. & Weil, D. P-Bodies: Cytosolic Droplets for Coordinated mRNA Storage. Trends Genet 34, 612–626 (2018).

63. Brothers, W.R., Ali, F., Kajjo, S. & Fabian, M.R. The EDC4-XRN1 interaction controls P-body dynamics to link mRNA decapping with decay. EMBO J 42, e113933 (2023).

64. Eulalio, A., Behm-Ansmant, I., Schweizer, D. & Izaurralde, E. P-body formation is a consequence, not the cause, of RNA-mediated gene silencing. Mol Cell Biol 27, 3970–81 (2007).

65. Luo, Y., Schofield, J.A., Simon, M.D. & Slavoff, S.A. Global Profiling of Cellular Substrates of Human Dcp2. Biochemistry 59, 4176–4188 (2020).

66. Banani, S.F. et al. Compositional Control of Phase-Separated Cellular Bodies. Cell 166, 651–663 (2016).

67. Kedersha, N., Tisdale, S., Hickman, T. & Anderson, P. Real-time and quantitative imaging of mammalian stress granules and processing bodies. Methods Enzymol 448, 521–52 (2008).

68. Poltronieri, P., Sun, B. & Mallardo, M. RNA Viruses: RNA Roles in Pathogenesis, Coreplication and Viral Load. Curr Genomics 16, 327–35 (2015).

69. den Boon, J.A., Diaz, A. & Ahlquist, P. Cytoplasmic viral replication complexes. Cell Host Microbe 8, 77–85 (2010).

70. Lavezzo, E. et al. G-quadruplex forming sequences in the genome of all known human viruses: A comprehensive guide. PLoS Comput Biol 14, e1006675 (2018).

71. Mendoza, O., Bourdoncle, A., Boule, J.B., Brosh, R.M., Jr. & Mergny, J.L. G-quadruplexes and helicases. Nucleic Acids Res 44, 1989–2006 (2016).

72. Pyle, A.M. Translocation and unwinding mechanisms of RNA and DNA helicases. Annu Rev Biophys 37, 317–36 (2008).

73. Heaton, S.M., Gorry, P.R. & Borg, N.A. DExD/H-box helicases in HIV-1 replication and their inhibition. Trends Microbiol 31, 393–404 (2023).

74. Tapescu, I. et al. The RNA helicase DDX39A binds a conserved structure in chikungunya virus RNA to control infection. Mol Cell 83, 4174–4189 e7 (2023).

75. Ribeiro de Almeida, C., et al. RNA Helicase DDX1 Converts RNA G-Quadruplex Structures into R-Loops to Promote IgH Class Switch Recombination. Mol Cell 70, 650–662 e8 (2018).

76. Wu, G., Xing, Z., Tran, E.J. & Yang, D. DDX5 helicase resolves G-quadruplex and is involved in MYC gene transcriptional activation. Proc Natl Acad Sci U S A 116, 20453–20461 (2019).

77. Shi, P. et al. SUMOylation of DDX39A Alters Binding and Export of Antiviral Transcripts to Control Innate Immunity. J Immunol 205, 168–180 (2020).

78. Rothan, H.A. et al. SARS-CoV-2 Infects Primary Neurons from Human ACE2 Expressing Mice and Upregulates Genes Involved in the Inflammatory and Necroptotic Pathways. Pathogens 11(2022).

79. Lee, B. et al. SARS-CoV-2 infection exacerbates the cellular pathology of Parkinson’s disease in human dopaminergic neurons and a mouse model. Cell Rep Med 5, 101570 (2024).

80. Yamazaki, T. et al. The closely related RNA helicases, UAP56 and URH49, preferentially form distinct mRNA export machineries and coordinately regulate mitotic progression. Mol Biol Cell 21, 2953–65 (2010).

81. Sugiura, T., Nagano, Y. & Noguchi, Y. DDX39, upregulated in lung squamous cell cancer, displays RNA helicase activities and promotes cancer cell growth. Cancer Biol Ther 6, 957–64 (2007).

82. Yoo, H.H. & Chung, I.K. Requirement of DDX39 DEAD box RNA helicase for genome integrity and telomere protection. Aging Cell 10, 557–71 (2011).

83. Xu, Z. et al. DDX39A resolves replication fork-associated RNA-DNA hybrids to balance fork protection and cleavage for genomic stability maintenance. Mol Cell 85, 490–505 e11 (2025).

84. Yellamaty, R. & Sharma, S. Critical Cellular Functions and Mechanisms of Action of the RNA Helicase UAP56. J Mol Biol 436, 168604 (2024).

85. Bi, H. et al. DDX41 resolves G-quadruplexes to maintain erythroid genome integrity and prevent cGAS-mediated cell death. Nat Commun 16, 7195 (2025).

86. Tandel, D. et al. SARS-CoV-2 Variant Delta Potently Suppresses Innate Immune Response and Evades Interferon-Activated Antiviral Responses in Human Colon Epithelial Cells. Microbiol Spectr 10, e0160422 (2022).

87. Qin, G. et al. RNA G-quadruplex formed in SARS-CoV-2 used for COVID-19 treatment in animal models. Cell Discov 8, 86 (2022).

88. Zhao, C. et al. Targeting RNA G-Quadruplex in SARS-CoV-2: A Promising Therapeutic Target for COVID-19? Angew Chem Int Ed Engl 60, 432–438 (2021).

89. Hondele, M. et al. DEAD-box ATPases are global regulators of phase-separated organelles. Nature 573, 144–148 (2019).

90. Marijan, D. et al. Protein thermal sensing regulates physiological amyloid aggregation. Nat Commun 15, 1222 (2024).

91. Gil-Garcia, M. et al. Local environment in biomolecular condensates modulates enzymatic activity across length scales. Nat Commun 15, 3322 (2024).

92. Lim, S. & Clark, D.S. Phase-separated biomolecular condensates for biocatalysis. Trends Biotechnol 42, 496–509 (2024).

93. Wegmann, S. et al. Tau protein liquid-liquid phase separation can initiate tau aggregation. EMBO J 37(2018).

94. Rupert, J., Monti, M., Zacco, E. & Tartaglia, G.G. RNA sequestration driven by amyloid formation: the alpha synuclein case. Nucleic Acids Res 51, 11466–11478 (2023).

95. Ahmad, S.R., AlShahrani, A.M., Vijayakumar, N. & Kumari, A. Pathogenic DDX39A Variant Disrupts Nuclear Homeostasis and Causes an Early-Onset Neurodegenerative Disorder With Cerebral Atrophy. Clin Genet (2025).

96. Zhang, S. et al. Post-translational modifications of soluble alpha-synuclein regulate the amplification of pathological alpha-synuclein. Nat Neurosci 26, 213–225 (2023).

97. Smeyne, R.J., Noyce, A.J., Byrne, M., Savica, R. & Marras, C. Infection and Risk of Parkinson’s Disease. J Parkinsons Dis 11, 31–43 (2021).

98. Marreiros, R. et al. Disruption of cellular proteostasis by H1N1 influenza A virus causes alpha-synuclein aggregation. Proc Natl Acad Sci U S A 117, 6741–6751 (2020).

99. Dadonaite, B. et al. The structure of the influenza A virus genome. Nat Microbiol 4, 1781–1789 (2019).

100. Ferhadian, D. et al. Structural and Functional Motifs in Influenza Virus RNAs. Front Microbiol 9, 559 (2018).

101. Meinhardt, J. et al. Olfactory transmucosal SARS-CoV-2 invasion as a port of central nervous system entry in individuals with COVID-19. Nat Neurosci 24, 168–175 (2021).

102. Yuan, X. et al. Propagation of pathologic alpha-synuclein from kidney to brain may contribute to Parkinson’s disease. Nat Neurosci 28, 577–588 (2025).

103. Kim, S. et al. Transneuronal Propagation of Pathologic alpha-Synuclein from the Gut to the Brain Models Parkinson’s Disease. Neuron 103, 627–641 e7 (2019).

104. Ahn, E.H. et al. Initiation of Parkinson’s disease from gut to brain by delta-secretase. Cell Res 30, 70–87 (2020).

105. Challis, C. et al. Gut-seeded alpha-synuclein fibrils promote gut dysfunction and brain pathology specifically in aged mice. Nat Neurosci 23, 327–336 (2020).

106. Holmqvist, S. et al. Direct evidence of Parkinson pathology spread from the gastrointestinal tract to the brain in rats. Acta Neuropathol 128, 805–20 (2014).

107. Arotcarena, M.L. et al. Bidirectional gut-to-brain and brain-to-gut propagation of synucleinopathy in non-human primates. Brain 143, 1462–1475 (2020).

108. Sharabi, Y., Vatine, G.D. & Ashkenazi, A. Parkinson’s disease outside the brain: targeting the autonomic nervous system. Lancet Neurol 20, 868–876 (2021).

109. Beatman, E.L. et al. Alpha-Synuclein Expression Restricts RNA Viral Infections in the Brain. J Virol 90, 2767–82 (2015).

110. Gupta, A. et al. Alpha-synuclein expression in neurons modulates Japanese encephalitis virus infection. J Virol 98, e0041824 (2024).

111. Kontopoulos, E., Parvin, J.D. & Feany, M.B. Alpha-synuclein acts in the nucleus to inhibit histone acetylation and promote neurotoxicity. Hum Mol Genet 15, 3012–23 (2006).

112. Koushik, S.V. & Vogel, S.S. Energy migration alters the fluorescence lifetime of Cerulean: implications for fluorescence lifetime imaging Forster resonance energy transfer measurements. J Biomed Opt 13, 031204 (2008).

113. Seibenhener, M.L. & Wooten, M.W. Isolation and culture of hippocampal neurons from prenatal mice. J Vis Exp (2012).

114. Volpicelli-Daley, L.A., Luk, K.C. & Lee, V.M. Addition of exogenous alpha-synuclein preformed fibrils to primary neuronal cultures to seed recruitment of endogenous alpha-synuclein to Lewy body and Lewy neurite-like aggregates. Nat Protoc 9, 2135–46 (2014).

115. Moutin, E. et al. Procedures for Culturing and Genetically Manipulating Murine Hippocampal Postnatal Neurons. Front Synaptic Neurosci 12, 19 (2020).

116. Hutchinson, E.C. et al. Mapping the phosphoproteome of influenza A and B viruses by mass spectrometry. PLoS Pathog 8, e1002993 (2012).

117. Rawat, S., Ghosh, S., Mondal, D., Anusha, V. & Raychaudhuri, S. Increased supraorganization of respiratory complexes is a dynamic multistep remodelling in response to proteostasis stress. J Cell Sci 133(2020).

118. Shevchenko, A., Wilm, M., Vorm, O. & Mann, M. Mass spectrometric sequencing of proteins silver-stained polyacrylamide gels. Anal Chem 68, 850–8 (1996).

119. Rappsilber, J., Ishihama, Y. & Mann, M. Stop and go extraction tips for matrix-assisted laser desorption/ionization, nanoelectrospray, and LC/MS sample pretreatment in proteomics. Anal Chem 75, 663–70 (2003).

120. Ge, S.X., Jung, D. & Yao, R. ShinyGO: a graphical gene-set enrichment tool for animals and plants. Bioinformatics 36, 2628–2629 (2020).

121. Caudron-Herger, M., Jansen, R.E., Wassmer, E. & Diederichs, S. RBP2GO: a comprehensive pan-species database on RNA-binding proteins, their interactions and functions. Nucleic Acids Res 49, D425–D436 (2021).

