## Supplementary Figures and Tables for "Cytosolic interaction with RNA-helicase DDX39A titrates viral RNA G-quadruplex mediated α-Synuclein amyloidogenesis"

Figure S1

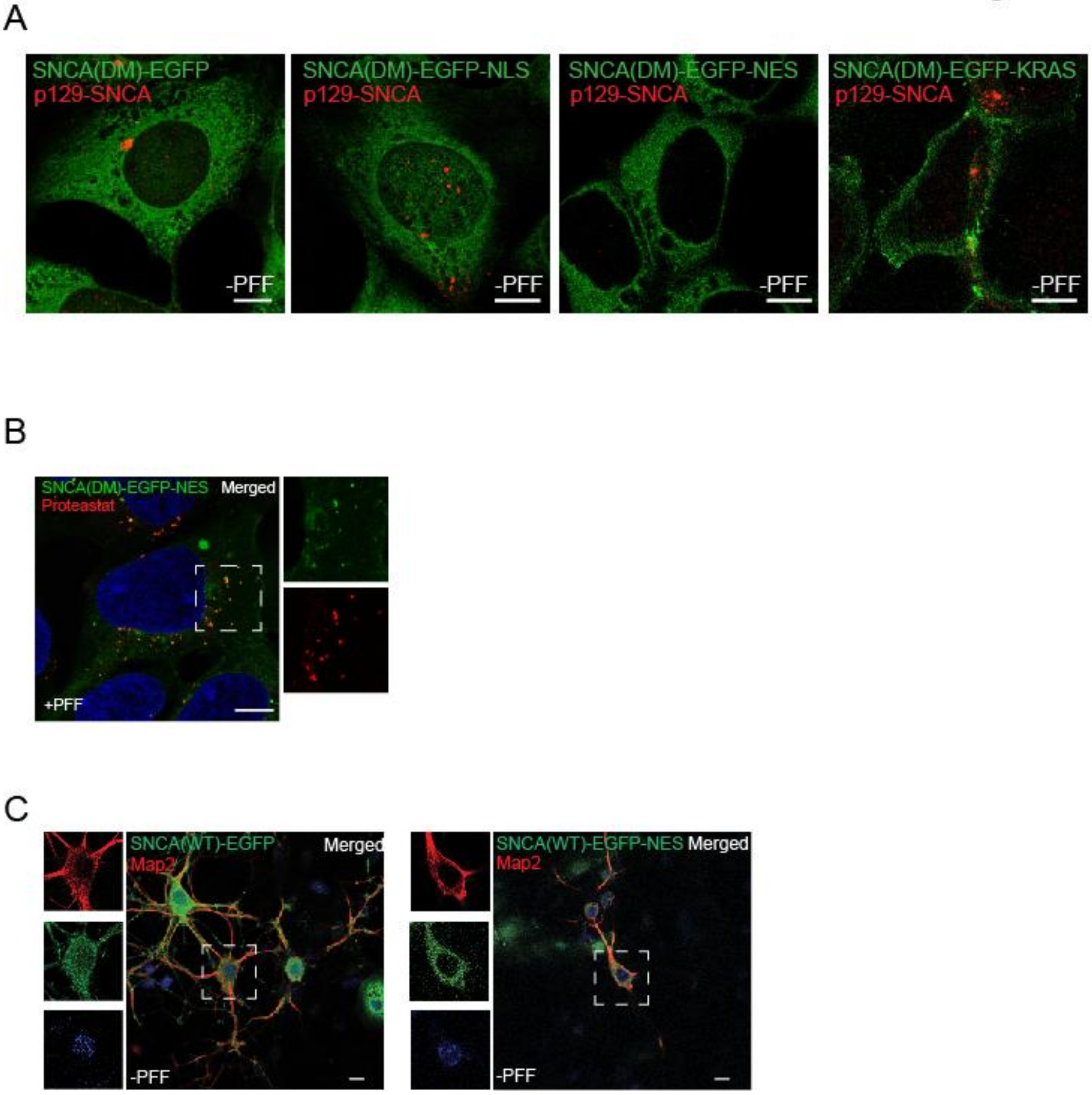

**Fig. S1: Subcellular location modulates α-Synuclein amyloidogenesis**

**A:** HEK293T stable lines expressing SNCA(DM)-EGFP and localization variants immunostained with p129-SNCA antibody (detecting only phosphorylated α-Synuclein (MJF-R13 (8-8)) and imaged.

**B:** HEK293T stable line expressing SNCA(DM)-EGFP-NES stained with ProteoStat dye. Insets shows single channel zoomed images of boxed regions.

**C:** Mice primary hippocampal neurons at DIV-6 were transduced with lentivirus with SNCA(WT)-EGFP or SNCA(WT)-EGFP-NES, treated with PFF at 13-DIV, fixed and immunostained with anti-MAP2 as neuronal marker at 20-DIV. Insets – single channel zoomed images of boxed regions.

Blue staining in microscopy images indicate nucleus (DAPI). Scale bar – 10 μm.

Figure S2

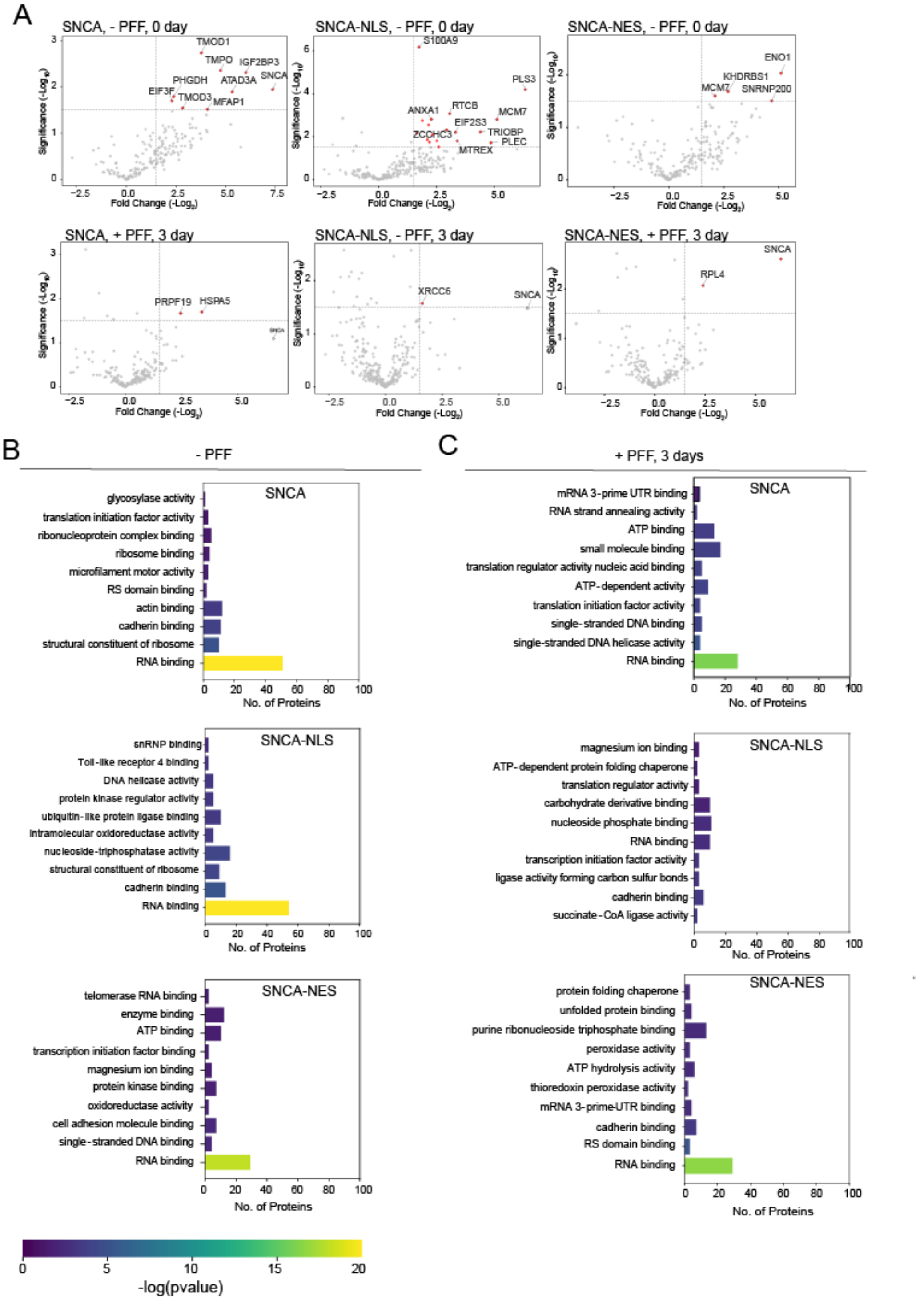

**Fig. S2: Spatiotemporal confinement modulates α-Synuclein interactions with RBPs.**
**A:** Volcano plots showing proteins interacting with α-Synuclein. The $-\log_{10}$ (p-value, One-sample *t*-test) plotted against the $\log_2$ fold change (SNCA-EGFP/EGFP, SNCA-EGFP-NLS/EGFP-NLS, and SNCA-EGFP-NES/EGFP-NES) of the abundance of the interacting proteins as identified by mass spectrometry experiments.

**B:** Gene Ontology (GO) enrichment analysis highlighting molecular functions significantly enriched for α-Synuclein interacting proteins in HEK293T stable line expressing SNCA(DM)-EGFP and localization variants in absence of PFF. FDR cut-off of 0.05 applied.
**C:** Gene Ontology (GO) enrichment analysis highlighting molecular functions significantly enriched for α-Synuclein interacting proteins in HEK293T stable line expressing SNCA(DM)-EGFP and localization variants after 3 days PFF-treatment. FDR cut-off of 0.05 applied.

Figure S3

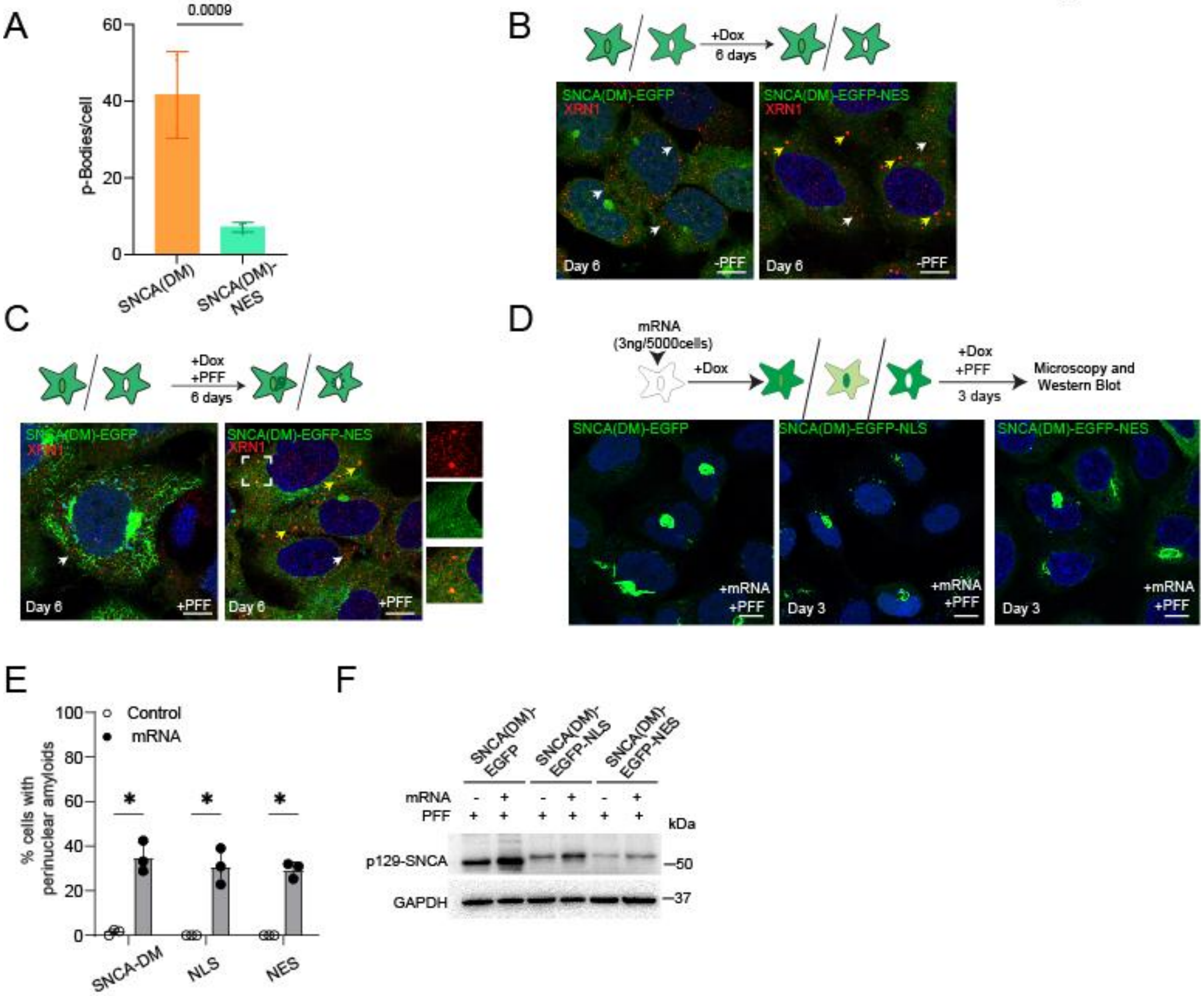

**Fig. S3: Cytosolic RNA-protein interactions modulate α-Synuclein amyloidogenesis**
**A:** Bar graph represents number of P-bodies in SNCA(DM)-EGFP and SNCA(DM)-EGFP-NES cells as shown in **3A**. N=4, ~25 cells counted in each biological repeats. Student t-test, Error bars indicate SD.
**B: Top -** Experimental Scheme. **Bottom**- Microscopy images showing SNCA(DM)-EGFP and SNCA(DM)-EGFP-NES cells in absence of PFF after 6 days, immunostained for XRN1.
**C: Top -** Experimental Scheme. **Bottom Left**- Microscopy images showing +PFF/±mRNA SNCA(DM)-EGFP-NES cells cultured for 3 days, immunostained for XRN1.
**D: Top -** Experimental Scheme. **Bottom** - Microscopy images showing +PFF/+mRNA HEK293T stable line expressing SNCA(DM)-EGFP and localization variants for 3 days.
**E:** Bar graph representing percentage of cells in **S3D** with perinuclear amyloids. N=3, Student t-test, Error bars represents SD. ~100 cells counted in each biological repeats.

**F:** Western blot for total cell lysates from **S3D** with p129-SNCA antibody - detecting only phosphorylated α-Synuclein (MJF-R13 (8-8)). GAPDH as loading control.
Blue staining in microscopy images indicate nucleus (DAPI). Scale bar – 10 μm.

Figure S4

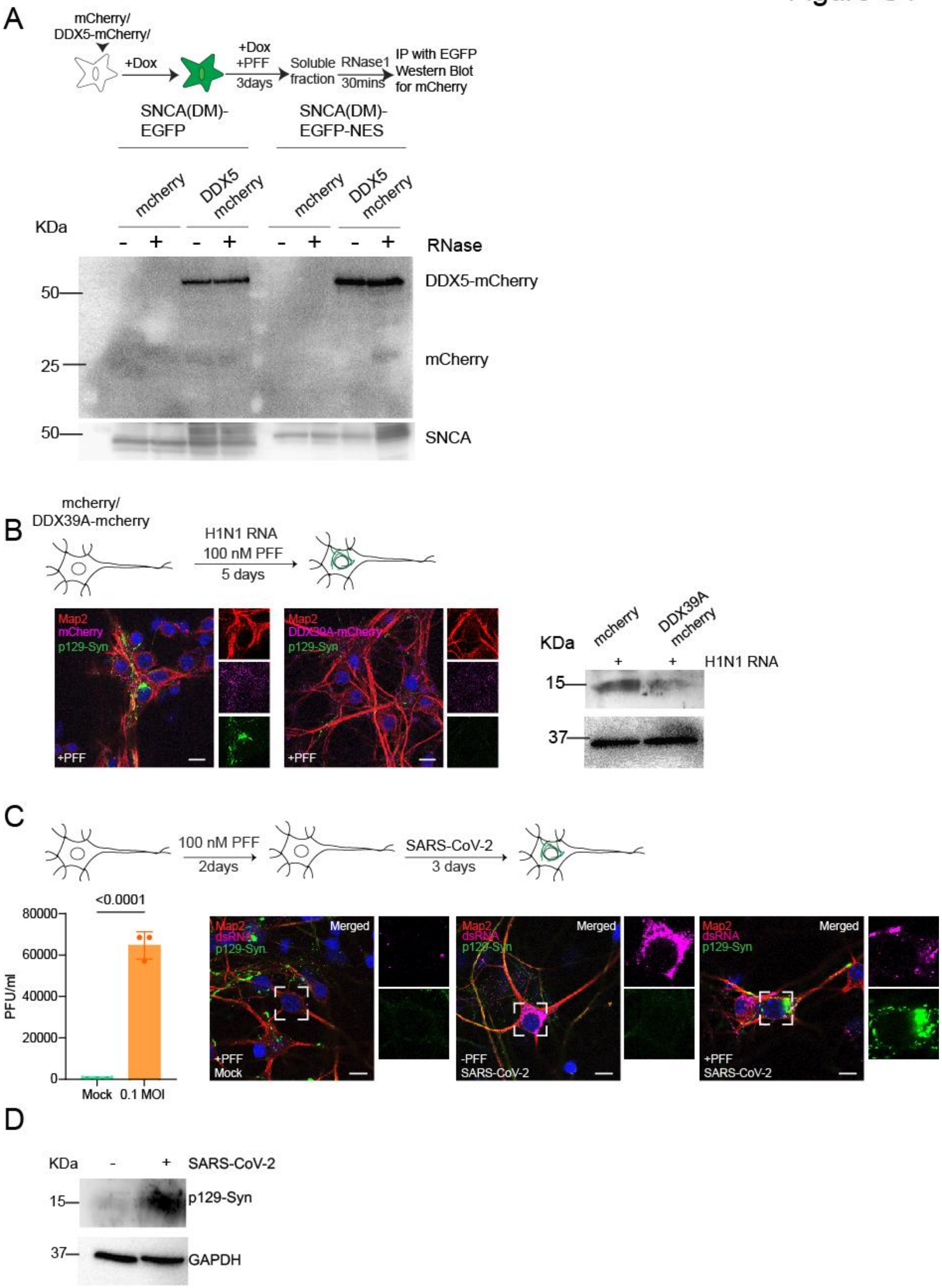

**Fig. S4: DDX39A alleviates rG4-mediated α-Synuclein amyloidogenesis.**
**A: Top -** Experimental scheme. **Bottom** – SNCA(DM)-EGFP and SNCA(DM)-EGFP-NES cells were transfected with mCherry and DDX5-mCherry and treated with PFF for 3 days. Immunoprecipitation (IP) for EGFP was performed with soluble fractions ±RNase1 (30 min) and probed with SNCA antibody - detecting both phosphorylated and non-phosphorylated α-Synuclein (14H2L1; epitope:117-125) and mCherry antibody.
**B: Top -** Experimental scheme. **Bottom Left** - Mice primary hippocampal neurons at DIV-5 were transfected with mCherry, DDX39A-mCherry with H1N1 genomic RNA were fixed after 4 days of PFF treatment and immunostained with anti Map2 antibody as neuronal marker and p129-SNCA antibody (detecting only phosphorylated α-Synuclein (MJF-R13 (8-8)) and imaged. Insets – zoomed single channel images. **Bottom Right** - Western Blot for total cell lysate of mice primary hippocampal neurons probed with p129-SNCA antibody - detecting only phosphorylated α-Synuclein (MJF-R13 (8-8)). GAPDH as loading control.
**C: Top -** Experimental scheme. **Bottom Left** – Virus titration of supernatant collected after 72 hrs of SARS-CoV-2 infected mice primary hippocampal neurons. **Bottom Righ**t – Mice primary hippocampal neurons at DIV-5 were treated with PFF. After 2 DIV neurons were transfected with mCherry, DDX39A-mCherry and DDX39ΔRNA-mCherry along with SARS-CoV-2 infection. Neurons were fixed after 3 days of infection and immunostained with anti Map2 antibody as neuronal marker, p129-SNCA antibody (detecting only phosphorylated α-Synuclein (MJF-R13 (8-8)), and viral dsRNA antibody; and imaged. Insets – zoomed single channel images.
**D:** Western Blot for total cell lysate of mice primary hippocampal neurons in **Fig. S4C** probed with p129-SNCA antibody - detecting only phosphorylated α-Synuclein (MJF-R13 (8-8)). GAPDH as loading control.
Error bars indicate SD from at least three independent experiments. Blue staining in microscopy images indicate nucleus (DAPI). Scale bar – 10 μm.

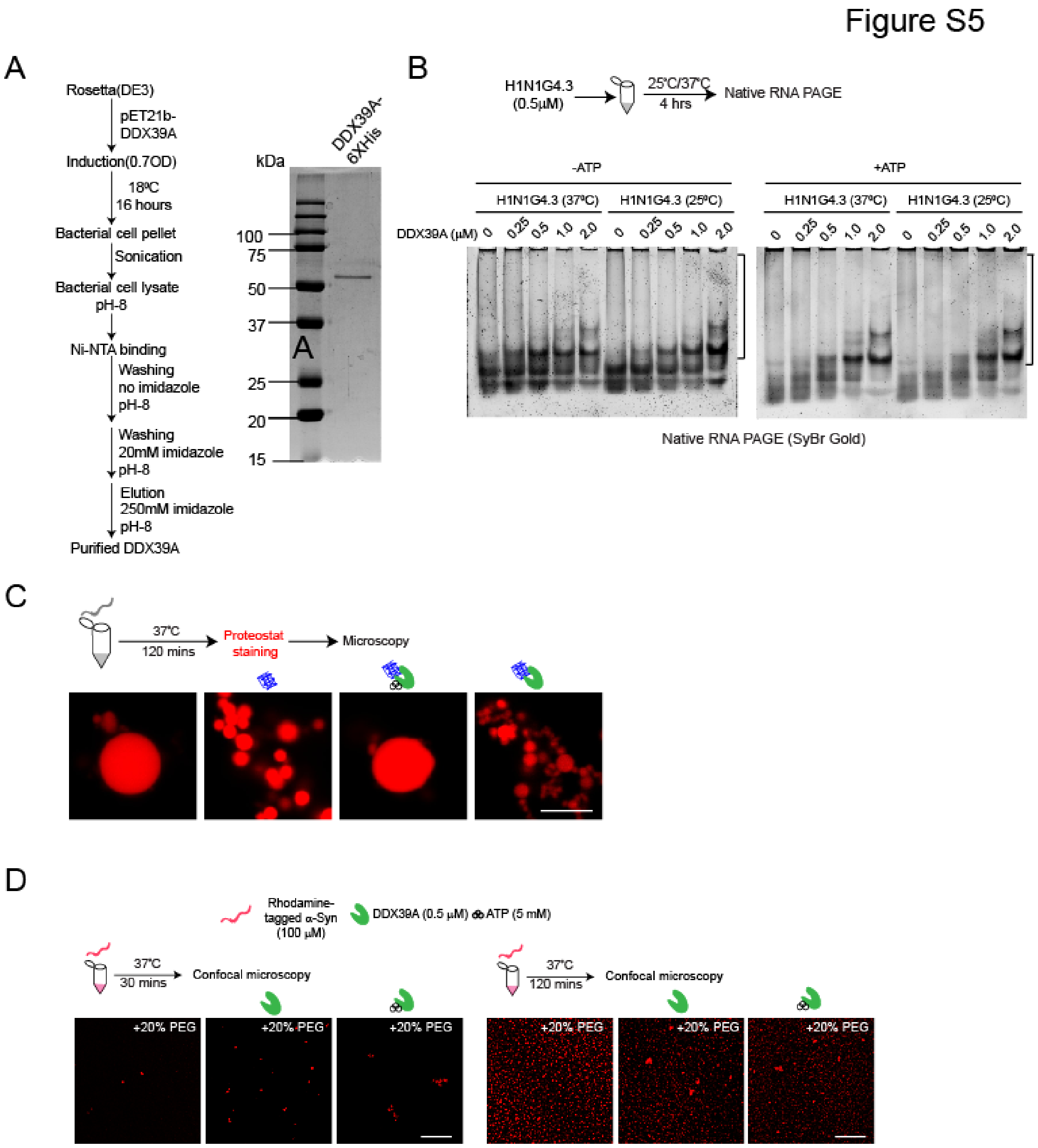

**Fig. S5: ATP dependent helicase activity of DDX39A prevents α-Synuclein sol-gel transition**

**A:** Purification protocol for DDX39A protein and coomassie blue stained SDS-PAGE.

**B: Top -** Experimental scheme. **Bottom -** Electrophoretic mobility shift assay (EMSA) for the interaction of DDX39A with H1N1G4.3 rG4 (500 nM) in absence (Left) and presence (Right) of 5 mM of ATP in a concentration dependent manner Brackets indicate H1N1G4.3-DDX39A complexes.

**C: Top -** Experimental scheme. **Bottom** - Microscopy images showing *In vitro* LLPS of purified recombinant human α-Synuclein (100 μM) in the presence of 20% PEG, after proteostat staining. Scale bar – 1 μm.

**D: Top -** Experimental scheme. **Bottom** - Microscopy images showing *in vitro* LLPS of purified recombinant rhodamine tagged human α-Synuclein (100 μM) in the presence of 20% PEG with and without 0.5 μM DDX39A and 5 mM ATP. Scale bar – 10 μm.

### Figure S6

A

| POSITION | SEQUENCE | G-SCORE | GENE |
|---|---|---|---|
| 313 | GGCUUUGGAGACUCCGUGGAGGAGG | 16 | ORF1ab-nsp1 |
| 604 | GGUAAUAAAGGAGCUGGUGG | 15 | ORF1ab-nsp1 |
| 1423 | GGUGGUCGCACUAUUGCCUUUGGAGG | 6 | ORF1ab-nsp2 |
| 1534 | GGUGUUGUUGGAGAAGGUUCCGAAGG | 18 | ORF1ab-nsp3 |
| 2674 | GGCGGUGCACCAACAAAGGUUACUUUUGG | 10 | ORF1ab-nsp3 |
| 3427 | GGAGGAGGUGUUGCAGG | 15 | ORF1ab-nsp3 |
| 4122 | GGUUAUACCUACUAAAAAGGCUGGUGG | 6 | ORF1ab-nsp3 |
| 4221 | GGGUUUAAAUGGUUACACUGUAGAGGAGG | 10 | ORF1ab-nsp3 |
| 8647 | GGAUACAAGGCUAUUGAUGGUGG | 14 | ORF1ab-nsp4 |
| 10221 | GGCUGGUAAUGUUCAACUCAGGGUUAUUGG | 9 | ORF1ab-3C-like proteinase |
| 13345 | GGUAUGUGGAAAGGUUAUGG | 19 | ORF1ab-nsp10 |
| 14907 | GGUUUUCCAUUUAAUAAAUGGGGUAAGG | 4 | Stemloop |
| 15168 | GGAACAAGCAAAUUCUAUGGUGGUUGG | 6 | Stemloop |
| 15408 | GGCGGUUCACUAUAUGUUAAACCAGGUGG | 3 | Stemloop |
| 18256 | GGAUUGGCUUCGAUGUCGAGGGG | 9 | Stemloop |
| 22276 | GGUGAUUCUUCUUCAGGUUGGACAGCUGG | 10 | S |
| 24175 | GGUUGGACCUUUGGUGCAGG | 17 | S |
| 24228 | GGCUUAUAGGUUUAAUGGUAUUGG | 19 | S |
| 25157 | GGCCAUGGUACAUUUGGCUAGG | 17 | S |
| 25911 | GGUGGUUAUACUGAAAAAUGGGAAUCUGG | 8 | ORF3a |
| 26706 | GGAUCACCGGUGGAAUUGCUAUCGCAAUGG | 7 | M |
| 28741 | GGCUUCUACGCAGAAGGGAGCAGAGGCGG | 9 | N |
| 28863 | GGCUGGCAAUGGCGG | 18 | N |
| 29083 | GGAAAUUUUGGGGACCAGG | 14 | N |
| 29194 | GGCAUGGAAGUCACACCUUCGGGAACGUGG | 11 | N |

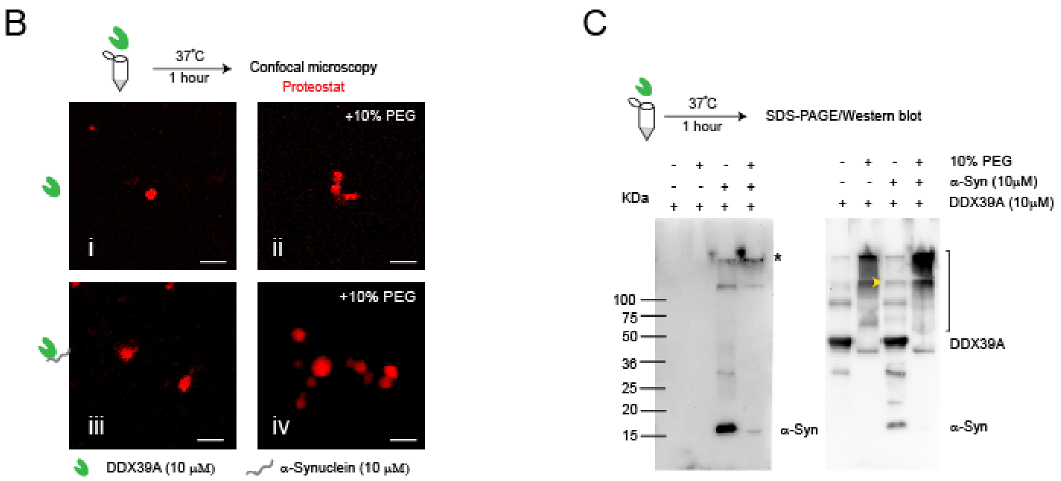

**Fig. S6: α-Synuclein facilitates DDX39A phase transition to enhance its helicase activity**

**A:** Table showing rG4 sequences in SARS-CoV-2 (B1.1.8) mapped in the genome, gene function and G-score as calculated by QRGS mapper.

**B: Top -** Experimental scheme. **Bottom** - Microscopy images showing *in vitro* LLPS of 10 µM purified DDX39A in absence and presence of 10% PEG with and without α-Synuclein after proteostat staining. Scale bar – 1µm.

**B: Top -** Experimental scheme. **Bottom –** Western blots of *in vitro* LLPS of 10µM purified DDX39A in the absence and presence of 10% PEG with and without 10µM α-Synuclein probed with anti- α-Synuclein antibody **(Left**) and anti-DDX39A antibody (**Right**).

### Methods Tables

| Methods Table S1 - Cloning Primers | | | |
|---|---|---|---|
| **Gene Name** | **Primer Sequence** | **Source** | **Catalog** |
| NLS | Forward – AGCTGTACAAGATGGCCGCAGATCCAAAAAA | Bioserve | NA |
| | Reverse – CCCTCTAGATTATACCTTTCTCTTCTTTTTTGGATC | Bioserve | NA |
| NES | Forward – AGCTGTACAAGATGAACCTGGTGGACCTCCA | Bioserve | NA |
| | Reverse – CCCTCTAGATTACTGCTGCTCGTCCAGC | Bioserve | NA |
| KRAS(SVIM) SDM | Forward –GAAAAAGAAGTCAAAGACAAAGAGTGTAATTATGTAA | Bioserve | NA |
| | Reverse – TTACATAATTACACTCTTTGTCTTTGACTTCTTTTTC | Bioserve | NA |
| SNCA(WT) (Lentivirus) | Forward – GGTACCGGTATGGATGTATTCATGAAAGGA | Bioserve | NA |
| | Reverse – CATCTCGAGGGCTTCAGGTTCGTAGTCTTGAT | Bioserve | NA |
| EGFP (Lentivirus) | Forward – AGCCCTCGAGATGGTGAGCAAGGGCGAGGAG | Bioserve | NA |
| | Reverse – TATCGAATTCCTTGTACAGCTCGTCCATGC | Bioserve | NA |
| EGFP-NES (Lentivirus) | Reverse – ATCGAATTCTTACTGCTGCTCGTCCAGCTC | Bioserve | NA |
| DDX39A | Forward - GCTTGGTACCATGGCAGAACAGGATGTGGAAAACG | Bioserve | NA |
| | Reverse - CCCTCTAGACCGGCTCTGCTCGATGTATGTG | Bioserve | NA |
| DDX5 | Forward - GCTTGGTACCATGTCGGGTTATTCGAGTGACCGAG | Bioserve | NA |
| | Reverse - CCGGCCGGTGGATCCTTTGGGAATATCCTGT | Bioserve | NA |

| | | | |
|---|---|---|---|
| mCherry | Forward - TAATCTAGAATGGTGAGCAAGGGCGAGGAGGATA | Bioserve | NA |
| | Reverse - GAAGGGCCCTTACTTGTACAGCTCGTCCATGCCGCC | Bioserve | NA |
| DDX39A(pET21 b) | Forward - GACAAGCTTATGGCAGAACAGGATGTGGAA | Bioserve | NA |
| | Reverse - GTGCTCGAGCCGGCTCTGCTCGATGTATGT | Bioserve | NA |

| **Methods Table S2 – Cell Number** | | |
|---|---|---|
| **Format** | **Media (ml)** | **Cell number** |
| 12-well plate (Nunc) | 1 | 5,000 |
| 6-well plate (Nunc) | 2 | 10,000 |
| 60mm dish (Nunc) | 3 | 15,000 |
| 150mm dish (Nunc) | 20 | 80,000 |

| **Methods Table S3 - Antibody List** | | | | |
|---|---|---|---|---|
| **Primary Antibody** | | | | |
| **Antibody Name** | **Company** | **Catalog** | **Dilution** | |
| | | | **ICC** | **Western** |
| Rabbit Monoclonal, alpha Synuclein Antibody (14H2L1) | Invitrogen | 701085 | 1:1000 | 1:10,000 |

| | | | | |
|---|---|---|---|---|
| Rabbit Monoclonal, Anti-Alpha-synuclein (phospho S129) Antibody [MJF-R13 (8-8)] | abcam | ab168381 | 1:1000 | 1:10,000 |
| Mouse Monoclonal, Anti-alpha synuclein (Phospho S129) Antibody (P-syn/81A) | abcam | ab184674 | 1:500 | NA |
| Anti-MAP2 antibody - Neuronal Marker | abcam | ab5392 | 1:1000 | NA |
| Anti-DCP1A (C-terminal) antibody produced in rabbit | sigma | D5454 | 1:100 | NA |
| Anti-XRN1 produced in rabbit, affinity isolated antibody | sigma | SAB4200028 | 1:500 | NA |
| Anti-RFP (RABBIT) Antibody | rockland | 600-401-379 | NA | 1:5000 |
| **Secondary Antibody** | | | | |
| Anti-Rabbit IgG (Whole molecule) (peroxidase conjugate) | sigma | A6154 | NA | 1:20,000 |
| Anti-mouse IgG (whole molecule) (peroxidase conjugate) | sigma | A4416 | NA | 1:20000 |
| Goat anti-Rabbit IgG (H+L) Highly Cross-Adsorbed Secondary Antibody, Alexa Fluor 488 | Invitrogen | A11034 | 1:1000 | NA |
| Goat anti-Mouse IgG (H+L) Cross-Adsorbed Secondary Antibody, Alexa Fluor 488 | Invitrogen | A11001 | 1:500 | NA |
| Goat anti-Rabbit IgG (H+L) Cross-Adsorbed Secondary Antibody, Alexa Fluor 647 | Invitrogen | A21244 | 1:1000 | NA |

| | | | | |
|---|---|---|---|---|
| Goat anti-Mouse IgG (H+L) Highly Cross-Adsorbed Secondary Antibody, Alexa Fluor Plus 647 | Invitrogen | A32728 | 1:500 | NA |
| Goat anti-Rabbit IgG (H+L) Cross-Adsorbed Secondary Antibody, Alexa Fluor 555 | Invitrogen | A21428 | 1:1000 | NA |
| Goat anti-Chicken IgY (H+L) Secondary Antibody, Alexa Fluor™ 647 | Invitrogen | A21449 | 1:1000 | NA |
